# Conformational dynamics and target-dependent myristoyl switch of calcineurin B homologous protein 3

**DOI:** 10.1101/2022.09.23.509142

**Authors:** Florian Becker, Simon Fuchs, Lukas Refisch, Friedel Drepper, Wolfgang Bildl, Uwe Schulte, Shuo Liang, Jonas Immanuel Heinicke, Sierra C. Hansen, Clemens Kreutz, Bettina Warscheid, Bernd Fakler, Evgeny V. Mymrikov, Carola Hunte

**Affiliations:** Institute of Biochemistry and Molecular Biology, ZBMZ, Faculty of Medicine, University of Freiburg, 79104 Freiburg, Germany; Faculty of Biology, University of Freiburg, 79104 Freiburg, Germany; Institute of Medical Biometry and Statistics (IMBI), Faculty of Medicine and Medical Center, University of Freiburg, 79104 Freiburg, Germany; Institute of Physics, University of Freiburg, 79104 Freiburg, Germany; Biochemistry and Functional Proteomics, Institute of Biology II, Faculty of Biology, University of Freiburg, 79104 Freiburg, Germany; Institute for Physiology, Faculty of Medicine, University of Freiburg, 79104 Freiburg, Germany; CIBSS - Centre for Integrative Biological Signalling Studies, University of Freiburg, 79104 Freiburg, Germany; Department of Biochemistry, Theodor-Boveri-Institute, University of Würzburg, 97074 Würzburg, Germany; BIOSS Centre for Biological Signalling Studies, University of Freiburg, 79104 Freiburg, Germany

**Author notes:** Correspondence: Evgeny Mymrikov and Carola Hunte, Institute of Biochemistry and Molecular Biology, ZBMZ, Faculty of Medicine, CIBSS, University of Freiburg, Stefan-Meier-Strasse 17, 79104 Freiburg, Germany. (E.M.) and (C.H.). contributed equally (Florian Becker, Simon Fuchs).

**Keywords:** CHP3, tescalcin, calcium ion, myristoylation, myristoyl switch, NHE1, conformational changes, target peptide, EF-hand Ca^2+^-binding protein

## Abstract

Calcineurin B homologous protein 3 (CHP3) is an EF-hand Ca^2+^-binding protein involved in regulation of cancerogenesis, cardiac hypertrophy and neuronal development via interactions with sodium/proton exchangers (NHEs) and signalling proteins. CHP3 binds Ca^2+^ with micromolar affinity providing the basis to respond to intracellular Ca^2+^ signals. Ca^2+^ binding and myristoylation are important for CHP3 function but the underlying molecular mechanism remained elusive. Here, we show that Ca^2+^ binding and myristoylation independently affect conformational dynamics and functions of human CHP3. Ca^2+^ binding increased flexibility and hydrophobicity of CHP3 indicative of an open conformation. CHP3 in open Ca^2+^-bound conformation had higher affinity for NHE1 and associated stronger with lipid membranes compared to the closed Mg^2+^-bound conformation. Myristoylation enhanced flexibility of CHP3 and decreased its affinity to NHE1 independently of the bound ion, but did not affect its binding to lipid membranes. The data exclude the proposed Ca^2+^-myristoyl switch for CHP3. Instead, they document a Ca^2+^-independent exposure of the myristoyl moiety induced by binding of the target peptide to CHP3 enhancing its association to lipid membranes. We name this novel regulatory mechanism “target-dependent myristoyl switch”. Taken together, the interplay of Ca^2+^ binding, myristoylation and target binding allows for a context-specific regulation of CHP3 functions.

## Introduction

The calcineurin B homologous protein 3 (CHP3, tescalcin) belongs to the EF-hand Ca^2+^-binding protein (EFCaBP) family (1) and is closely related to calcineurin B homologous proteins CHP1 and CHP2 (2). CHPs interact with several isoforms of sodium/proton exchangers (NHEs), and this interaction is required for the localization and function of NHE transporters on the plasma membrane (3). Besides NHEs, CHP3 interacts with calcineurin A (4), subunit 4 of COP9 signalosome (5) and glycogen synthase kinase 3 (GSK3) (6). Recent studies revealed that the CHP3 expression level correlates with the progression, metastasis and invasiveness of gastric, renal, and colorectal cancers (7-10). In addition, genome-wide association studies in combination with neuroimaging identified CHP3 as a key regulator of neurogenesis for hippocampal volume formation (11-13). Via regulation of GSK3 and calcineurin activities, it seems to counteract cardiac hypertrophy (6, 14). Thus, CHP3 is an emerging important player in cellular Ca^2+^ signalling networks involved in the regulation of cell proliferation and development in different tissues. The underlying molecular mechanisms are not well understood.

All three CHPs were shown to undergo conformational changes upon Ca^2+^ binding (4, 15). Notably, CHP3 has a lower affinity for Ca^2+^ (0.8 µM) (4) in comparison to CHP1 and CHP2 (K_D_ values below 100 nM) (16, 17). This may enable CHP3 to respond to Ca^2+^ signals with conformational changes when the intracellular Ca^2+^ concentration elevates from 100 nM to 1 µM or higher (18). Ca^2+^-induced conformational changes are characteristic for Ca^2+^-sensor proteins such as calmodulin (CaM) and calcineurin B (19, 20). At the resting Ca^2+^ level, they adopt a closed conformation, in which the hydrophobic target binding pocket is occluded. Upon a rise of the intracellular Ca^2+^ concentration, Ca^2+^ binding to EF-hand(s) causes an “opening” of this pocket providing the structural basis for the transmission of Ca^2+^ signals (20). This opening can trigger the binding of EFCaBPs to their target proteins (21). In addition, Ca^2+^-induced conformational changes of EFCaBPs such as calcineurin B or GCAPs stably associated with target proteins can affect the function of the latter (19, 22). Recently, we demonstrated an increase of CHP3 hydrophobicity upon Ca^2+^ binding that most likely resulted from the opening of the hydrophobic target binding pocket (15). CHP3 binds also Mg^2+^ with a low affinity (*K*_D_ = 73 µM) in the absence of Ca^2+^. The presence of Mg^2+^ reduces the affinity for Ca^2+^ from 0.8 µM to 3.0 µM, indicating a direct competition between Ca^2+^ and Mg^2+^ for the EF-3 binding site (4). Thus, CHP3 should be present in the Mg^2+^-bound state in a cell at basal Ca^2+^ concentration, ready to respond to Ca^2+^ signals, yet, the exact mechanism is not fully described.

In addition, CHP3 harbours an N-terminal myristoylation site similar to other EFCaBPs such as recoverin and calcineurin B (4, 23). This modification often enhances protein binding to cellular membranes, but it can also stabilize the structure of a protein and/or regulate its function (24, 25). Membrane binding via the myristoyl moiety is usually enforced with either a second lipidation site or clusters of positively charged and/or hydrophobic residues (24, 25). The exposure of the myristoyl moiety from the modified protein and thereby its binding to lipid membranes can be regulated by various signals, for instance by exchange of GDP to GTP (ligand-dependent myristoyl switch in ADP ribosylation factor 1 (ARF1) GTPase) (26), by phosphorylation (phosphorylation myristoyl switch in myristoylated alanine-rich C kinase substrate (MARCKS) (27) and in the C-subunit of protein kinase A (28)) or by pH change (pH-dependent myristoyl-histidine switch in hisactophilin (29)). In many Ca^2+^-sensor proteins (for instance recoverin, neurocalcin, visinin-like proteins), the myristoyl group becomes accessible to lipid membranes after Ca^2+^ binding (22). This mechanism is called Ca^2+^-myristoyl switch. However, not all myristoylated EFCaBPs have a Ca^2+^-myristoyl switch. In the neuronal calcium sensor-1 (NCS-1), the myristoyl moiety becomes exposed in the presence of lipid membranes even at low Ca^2+^ concentration (22, 30, 31); and it is constantly hidden within the protein core in guanylyl cyclase activating proteins (GCAPs) (22). Myristoylated CHP1 binds to microsomal membranes in a Ca^2+^-dependent manner indicating the presence of a Ca^2+^-myristoyl switch (32), whereas binding of myristoylated CHP3 to lipid membranes has not been reported so far. Co-expression of NHE1 with CHP3 that lacked myristoylation and/or Ca^2+^-binding sites significantly reduced the half-life at the cell surface and the activity of this transporter (23). Simultaneous myristoylation and Ca^2+^-binding was suggested to be important for NHE1 stabilization by CHP3 (23). Based on these results, the presence of a Ca^2+^-myristoyl switch for CHP3 was proposed (4, 23), though the exposure of the myristoyl group in response to Ca^2+^ binding has not been shown experimentally.

The interaction of CHP3 with NHE1 is an ideal system to analyse the effects of myristoylation and Ca^2+^ binding on CHP3. We previously demonstrated that CHP3 binds with high affinity to the CHP-binding region of human NHE1 (CBD) in the presence of Mg^2+^ (33), and it was shown by co-immunoprecipitation of CHP3-myc and NHE1-HA that addition of Ca^2+^ increased the amount of the complex formed (23). Here, we show that Ca^2+^ and N-terminal myristoylation independently regulate conformational dynamics of CHP3 and its interaction with NHE1 providing the molecular basis for regulation of CHP3 function. This excludes a Ca^2+^-myristoyl switch in CHP3, instead, surface exposure of the myristoyl moiety was triggered by target peptide binding as probed by interaction with liposomes. We named this novel mechanism “target-dependent myristoyl switch”. Our study provides fundamental mechanistic understanding of the regulation of CHP3 function.

## Results

### Pure fully myristoylated and non-myristoylated untagged CHP3 are functional in binding a single calcium ion

In order to dissect the effects of Ca^2+^ and myristoylation on target and lipid binding of human CHP3, we aimed for pure untagged CHP3 and myristoylated CHP3 (mCHP3). Non-myristoylated CHPs were previously produced with affinity tags and purified by corresponding affinity chromatography (4, 15, 33). Yet, the peptide tag used for affinity purification of rat CHP1 was shown to interact with the hydrophobic target binding pocket (34) and might interfere with conformational changes. Myristoylated untagged CHP1 had been produced with low yield by co-expression with yeast N-myristoyltransferase (35). Here, we produced untagged CHP3 and mCHP3, the latter by co-expression with human N-myristoyl transferase 1, and used Ca^2+^-dependent hydrophobic interaction chromatography (HIC) for purification. HIC has been previously used for purification of other EFCaBPs including calmodulin (CaM) and calcineurin B (36, 37). Combining HIC and gel filtration, we obtained pure untagged proteins, as shown by SDS-PAGE analysis (Figure 1A) with an average yield of 9 mg per liter of expression culture. Single symmetrical peaks in the elution profiles of analytical gel filtration indicated monodisperse protein preparations of CHP3 and mCHP3. Both proteins form only monomers under reducing conditions in the presence of 2 mM TCEP (Figure S1). The slightly higher electrophoretic mobility of mCHP3 compared to CHP3 resolved in high-resolution SDS-PAGE analysis already indicated that the protein was covalently modified (Figure 1A). To check the degree of myristoylation, we analysed mCHP3 and CHP3 by native mass spectrometry (MS). The measured mass of mCHP3 was increased by ∼211 Da (Figure 1B, bottom) in comparison to CHP3 (Figure 1B, top), which is in agreement with the covalent attachment of one myristoyl group (M_r_=210 Da). Native MS analysis of three independently produced and purified samples confirmed reproducible complete myristoylation of recombinant CHP3. Interestingly, the ion mobility arrival time was shorter for mCHP3 (Figure 1C) indicating a more compact shape of mCHP3 compared to CHP3. To reveal the stoichiometry of Ca^2+^-binding, we performed native MS analysis of CHP3 and mCHP3 after addition of Ca^2+^. The molecular masses of both proteins were shifted by ∼40 Da (Figure 1B), which corresponds to the binding of a single Ca^2+^ ion and, thus, documents that both recombinant CHP3 and mCHP3 are functional in respect to Ca^2+^ binding and confirms the presence of one Ca^2+^-binding site (4, 38).

**Figure 1.**
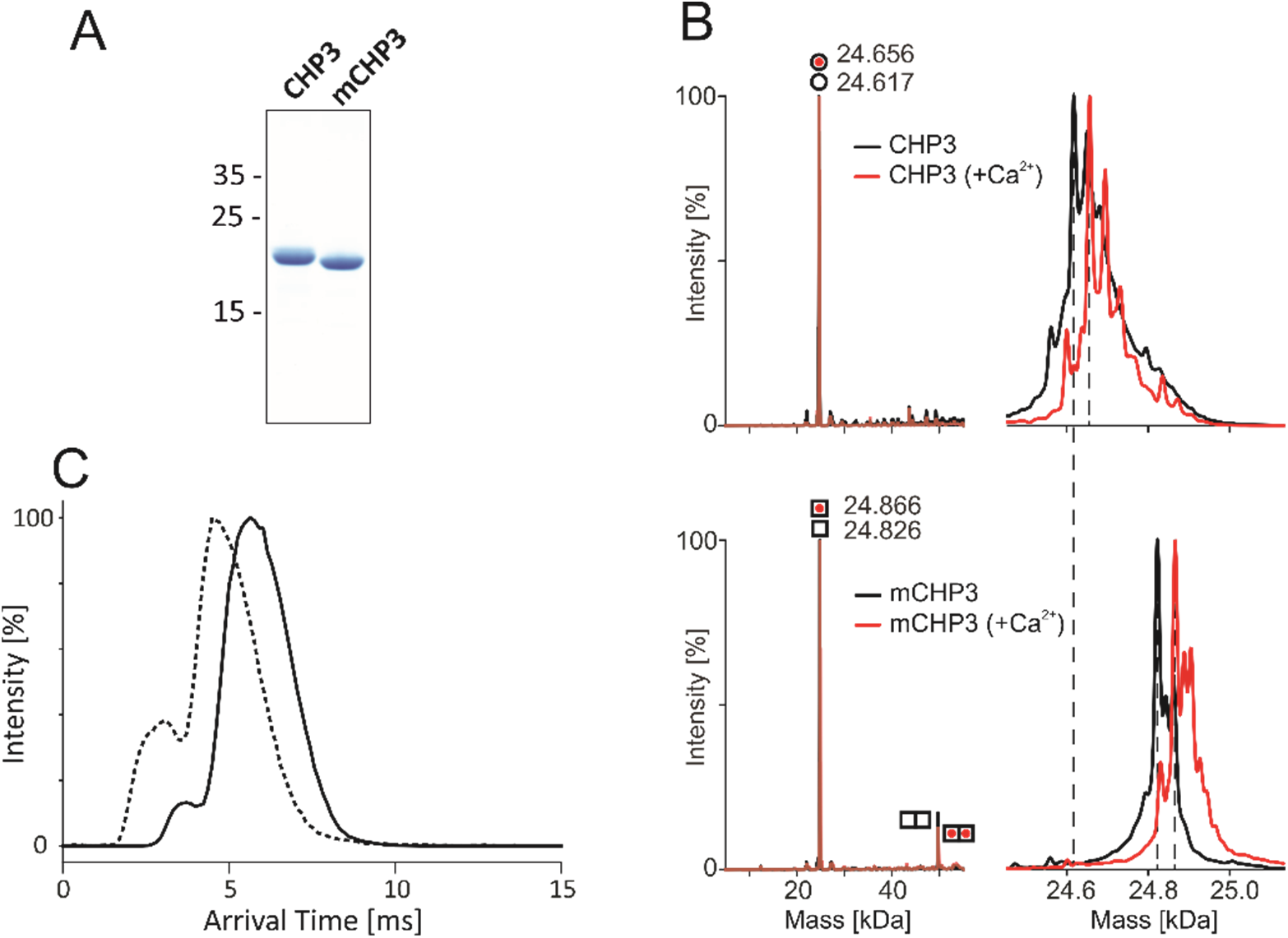
Quality control of the purified non-myristoylated CHP3 and myristoylated CHP3 (mCHP3). (**A**) SDS-PAGE analysis (16%, tricine mini gels) showed a mobility shift of mCHP3 in comparison to CHP3; positions of co-migrated molecular mass standards are indicated in kDa on the left. (**B**) Left, deconvoluted mass spectra of CHP3 (top) and mCHP3 (bottom) show a measured intact protein mass of 24.617 kDa and 24.828 kDa (ΔM_r_=211 Da), which is in line with the addition of the myristoyl group of 210 Da. For mCHP3, also low abundant dimeric species were observed by native MS analysis. Right, zoom-in spectra in the range of 24.8 ± 0.3 kDa that show differences in masses as indicated by dashed lines. In the presence of Ca^2+^, the measured intact protein mass of CHP3 and mCHP3 was increased by ∼40 Da (mass accuracy ± 1Da), respectively, indicating binding of one Ca^2+^ ion. (**C**) Ion mobility arrival time distributions for the +8 charge state of CHP3 (solid line) and mCHP3 (dashed line). Arrival times of mCHP3 were reduced by 1.36 ms (± 0.37 ms; n=3; p=0.024) compared to CHP3. See Figure S2 for non-deconvoluted mass spectra showing charge state distributions of CHP3 and mCHP3 measured in the positive ion mode by native MS.

### Ca^2+^-induced conformational changes are similar in CHP3 and mCHP3

Next, we analysed the effect of Ca^2+^ binding in the presence of Mg^2+^ and of the N-terminal myristoylation on CHP3 conformation. For this purpose, we optimized the FPH (fluorescence probe hydrophobicity) assay, previously developed to monitor Ca^2+^-induced conformational changes of CHPs (15), by using the dye ProteOrange (Lumiprobe) at defined micromolar concentration (see Materials and Methods). In this assay, the fluorescence of the dye strongly increases upon its binding to hydrophobic protein surfaces (39).

In the Mg^2+^-bound state, CHP3 and mCHP3 showed low fluorescence (Figure S3A). After Ca^2+^ addition, the fluorescence intensity increased by 30% and 40% for CHP3 and mCHP3, respectively, reflecting an increase in hydrophobicity (Figure 2A). Ca^2+^ removal by EGTA reduced the fluorescence to the initial level, indicating the reversibility of Ca^2+^ binding and of the respective conformational changes. Depletion of both Mg^2+^ and Ca^2+^ by EDTA turned the protein into the non-physiological apo-state, the fluorescence was in between that of the closed Mg^2+^-bound and open Ca^2+^-bound states. The proteins appeared to be destabilized in the apo-state and could not be reverted into the functional state by Ca^2+^ addition (Figure 2A). Interestingly, the N-terminal myristoylation did not affect the conformational changes of CHP3. As a control, we probed the hydrophobicity of recoverin, the prototypic protein with a classical Ca^2+^-myristoyl switch (Figure S3B). Ca^2+^ induced similar changes of non-myristoylated recoverin as of CHP3 in the FPH assay, whereas hydrophobicity of the myristoylated protein increased much stronger (about 3.5 fold) in response to Ca^2+^ binding, which most likely resulted from the exposure of the myristic group.

**Figure 2.**
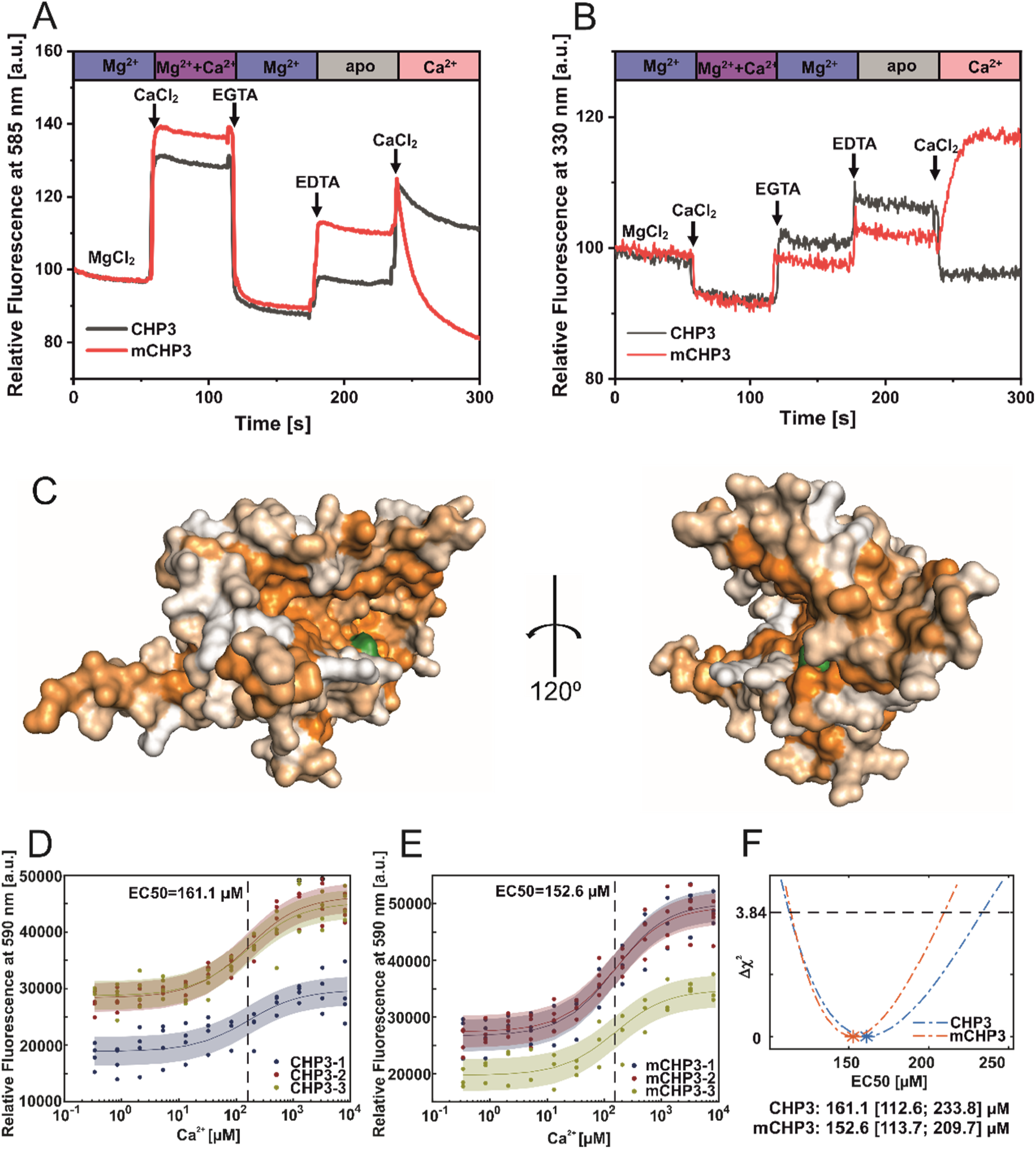
Ca^2+^-induced conformational changes in CHP3 and mCHP3. (**A**) Kinetic fluorescence probe hydrophobicity (FPH) assay. Fluorescence of dye bound at hydrophobic protein surfaces was monitored at λ_em_=585 nm (excitation λ_ex_=470 nm) and at 22°C. Protein (1.5 µM) was prepared in the Mg^2+^-bound state (2 mM MgCl_2_, 1 mM EGTA), and CaCl_2_, EDTA and EGTA were sequentially added as indicated. First, 2 mM CaCl_2_ was added and then chelated with addition of 3 mM EGTA. Next, 3 mM EDTA was added to remove both divalent ions (CHP3 in apo-state) followed by another addition of 4 mM CaCl_2_. (**B**) Intrinsic tryptophan fluorescence was monitored at λ_em_=330 nm (excitation λ_ex_=280 nm) at 22°C. Protein (2.5 μM) in the Mg^2+^-bound state was used and the additions were performed as described above for the FPH assay. (**C**) AlphaFold model (40) of CHP3 in surface presentation, with surface coloured for hydrophobicity (64). The model most likely resembles the open or target-bound conformation. The single tryptophan residue highlighted in green (Trp191) is located in the hydrophobic target binding pocket. (**D-F**) EC50 values for binding of Ca^2+^ to CHP3 (**D**) and mCHP3 (**E**) determined with FPH assay. Fluorescence of samples with CHP3 or mCHP3 at different Ca^2+^ concentrations was measured at 590 nm in the presence of 2 mM MgCl_2_. Three biological replicates (shown in different colours) with 3-4 technical replicates each for CHP3 and mCHP3 were measured. The data were fitted with Hill equation using global non-linear regression. (**F)** 95% confidence intervals (Δχ2 of 3.84) of EC50 values for binding of Ca^2+^ calculated with profile likelihood method. EC50 values (asterisks) are shown with confidence intervals in square brackets below the graph.

To prove that changes in the dye-mediated fluorescence observed in the FPH assay were indeed caused by conformational changes, we probed the intrinsic tryptophan fluorescence of CHP3 and mCHP3. According to the 3D model of CHP3 predicted with AlphaFold2.0 (40), the single tryptophan residue is located in the hydrophobic pocket of CHP3 (Figure 2C). In general, intrinsic tryptophan fluorescence increases in a hydrophobic environment, whereas an exposure of a tryptophan residue to the solvent quenches its fluorescence (41). The observed decrease of the intrinsic fluorescence after Ca^2+^ addition is in line with “opening” of the hydrophobic pocket (Figure 2B) and fully correlates with the increase in CHP3 hydrophobicity observed with the FPH assay. These conformational changes monitored by intrinsic fluorescence were also reverted by Ca^2+^ removal (EGTA addition) (Figure 2B). Only for mCHP3, the removal of both Mg^2+^ and Ca^2+^ caused irreversible changes in the intrinsic fluorescence.

In order to evaluate whether myristoylation affects the Ca^2+^ binding affinity of CHP3, we determined the EC50 values for binding of Ca^2+^ to CHP3 and mCHP3 in the presence of Mg^2+^ using the FPH assay. Addition of Ca^2+^ at saturating concentration increased the fluorescence (Figure 2A). We now titrated Ca^2+^ concentration from 0.3 µM to 8 mM and measured fluorescence using biological and technical replicates (Figure 2D,E). We determined Ca^2+^ EC50 values performing a global fit for all data. The difference between EC50’s obtained for CHP3 (161.1 [112.6; 233.8] μM; Figure 2D) and mCHP3 (152.6 [113.7; 209.7] µM; Figure 2E) is insignificant, indicating that myristoylation does not affect the Ca^2+^-binding properties of CHP3.

Combining the data of the FPH assay and of the intrinsic tryptophan fluorescence, we conclude that CHP3 undergoes reversible Ca^2+^-induced conformational changes as typical for a Ca^2+^-sensor protein. The hydrophobic target binding pocket of CHP3 is occluded in the Mg^2+^-bound (closed) state, and becomes exposed to the environment upon Ca^2+^ binding (open state).

### Ca^2+^ binding and myristoylation independently affect thermal stabilities of CHP3 and its complex with NHE1 target peptide

To further dissect the effects of Ca^2+^ and N-terminal myristoylation on the dynamic nature of CHP3, we probed the thermal stability of CHP3 alone and in complex with the CHP-binding domain of NHE1 (CBD, NHE1 residues 525-545) using nano-differential scanning fluorimetry (nanoDSF). To obtain the CHP3:CBD complex used for nanoDSF, we co-expressed CHP3 and CBDHis using the pETDuet-1 system and purified the resulting complex. For the myristoylated complex (mCHP3:CBD), simultaneous co-expression of three proteins (CHP3, CBDHis and human N-myristoyl transferase 1 (NMT1)) from a modified pETDuet-1 vector was performed, and the resulting complex was purified. Myristoylation of CHP3 in the complex was confirmed by ESI-TOF mass spectrometric analysis. Free CHP3 had the highest thermal stability in the closed Mg^2+^-bound state (T_m_ 70.1±0.3°C), and Ca^2+^ binding reduced the melting point to 66.9±0.2°C (CHP3 in Figure 3). In the presence of both ions, the Ca^2+^ effect appeared to be dominant with a melting temperature (T_m_) of 65.0±0.9°C. In line with the results of the FPH assay which indicated a destabilized apo-state, the latter showed strongly reduced thermal stability (T_m_ 56.1±0.5°C). Notably, binding of the target peptide CBD strongly increased the thermal stability for the Ca^2+^-bound and apo-states (compare CHP3 and CHP3:CBD in Figure 3), with about 12°C in the presence of Ca^2+^, 13°C in the presence of Ca^2+^ plus Mg^2+^ and about 11°C for the apo-state. Thus, the CHP3:CBD complex showed the highest thermal stability when Ca^2+^ was bound. Surprisingly, CBD binding did not have any effect on the thermal stability of the Mg^2+^-bound state.

**Figure 3.**
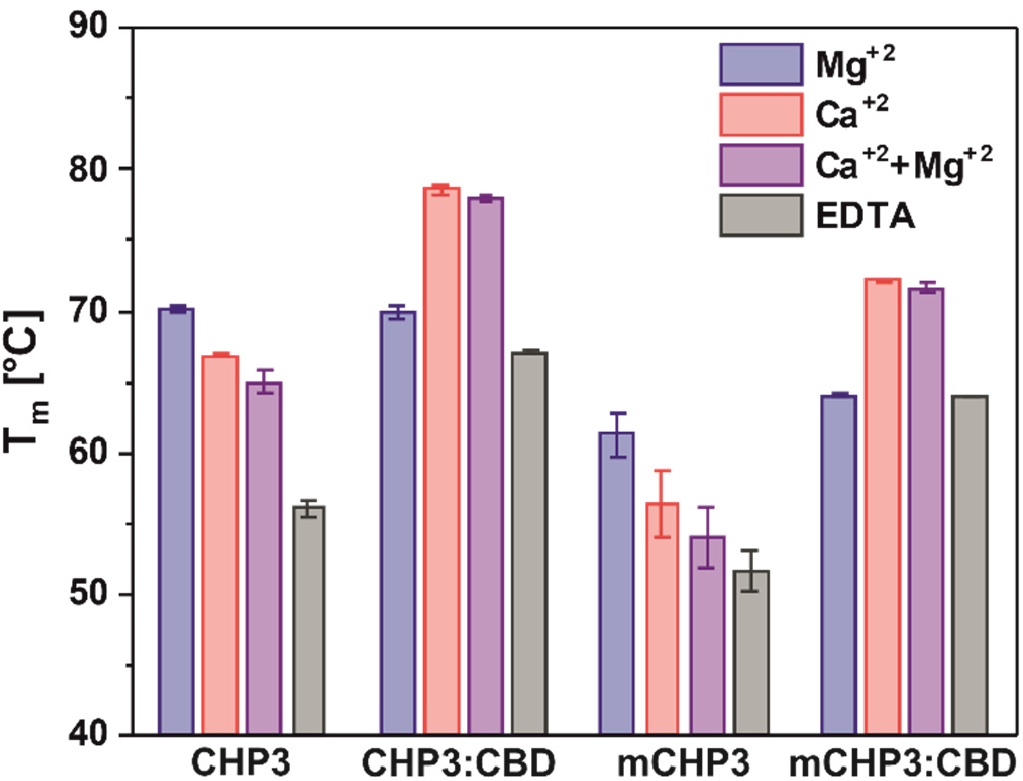
Ca^2+^ and target peptide binding affect the thermal stability of CHP3 and mCHP3. Thermal stabilities of free proteins and complexes with the target peptide CBD were measured with nanoDSF in the presence of either 10 mM Mg^2+^, Ca^2+^, Ca^2+^ + Mg^2+^ or EDTA. Temperatures of thermal unfolding (T_m_) are shown as mean±SD of three independent biological replicates. Raw nanoDSF traces are shown in Figure S4.

The N-terminal myristoylation decreased the T_m_ of CHP3 in all conformational states to the same degree (9.8°C for Mg^2+^-bound and 10.6°C for Ca^2+^-bound states, 10.9°C in the presence of both ions, 4.5°C for the already destabilized apo-state) (compare CHP3 and mCHP3 in Figure 3). A similar destabilizing effect of the myristoylation was observed for the mCHP3:CBD complex with a decrease of T_m_ in the range from 6.4°C for the Ca^2+^-bound state to 3.2°C for the apo-state (compare CHP3:CBD and mCHP3:CBD in Figure 3). Thus, the N-terminal myristoylation affects the conformation and/or flexibility of CHP3 independently of the bound ion and of target binding.

### Ca^2+^ binding to CHP3 allows more effective cleavage within EF-2, whereas CBD binding completely prevents tryptic digestion

In order to pinpoint areas with increased protein flexibility in a given conformational state and to identify regions affected by ion binding and myristoylation, we analysed the proteolytic stability of CHP3 in a state-dependent manner. We performed limited trypsinolysis of CHP3 and mCHP3 and of their complexes with CBD in the presence of Ca^2+^, Mg^2+^ or EDTA.

CHP3 was readily proteolysed by trypsin with nearly one half of the protein already degraded after 5 minutes and almost no full-length protein remained after 60 min incubation (Figure 4A, top (FL). During the reaction, we observed the appearance of two major fragments (labelled 1 and 2 in Figure 4A). We analyzed those fragments with mass spectrometry to determine exact masses and precisely locate the cleavage sites (Figure S5B). Fragment 1 has a mass of 17.92 kDa corresponding to residues 2-155 of CHP3 (UniProtID Q96BS2-1) lacking the C-terminal part (Figure 4C). Fragment 2 has a mass of 9.63 kDa and includes residues 74-155. Thus, it derived from a cleavage of fragment 1 within the predicted EF-2 (Figure 4C). Noteworthy, both fragments still contain the single functional EF**-**hand of CHP3 (EF-3). Target peptide binding nearly completely prevented the trypsin cleavage, and the intensity of the full-length CHP3 band was only slightly reduced even after 60 min incubation (Figure 4A, bottom). Next, we compared the proteolytic stability of CHP3 in different conformational states. In both ion-bound states as well as in the presence of both ions, the degradation rate of the full-length protein (FL) was comparable, whereas the apo-state degraded much faster (Figure 4B, FL). Fragment 1 appeared already after 5 min and degraded further over time (Figure 4B, 1), its cleavage was more pronounced in the presence of Ca^2+^ and was accompanied with an increase of fragment 2 (Figure 4B, 2). This indicates that the cleavage site located in EF-2 becomes slightly more accessible in the open Ca^2+^-bound state. N-terminal myristoylation did not affect the proteolytic stability of the full-length protein, and just slightly reduced the stability of fragment 1 in all conformational states (Figure 4B). The complex formation with CBD shielded both CHP3 and mCHP3 from the tryptic cleavage, nearly no degradation occurred (Figure 4B, CHP3:CBD and Figure S5A).

**Figure 4.**
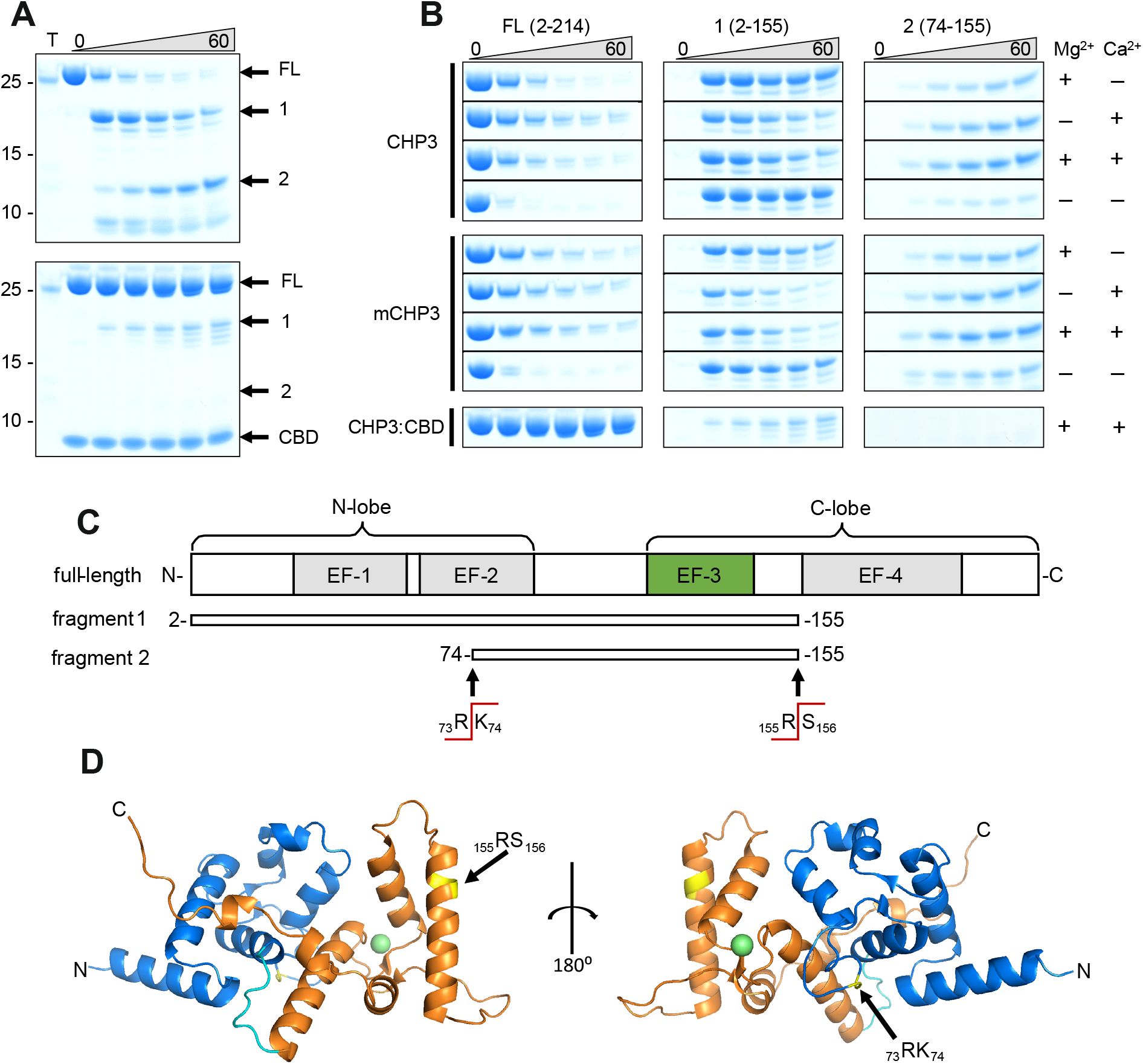
Ca^2+^- binding and complex formation change the accessibility of trypsin cleavage sites in CHP3 and mCHP3. (**A**) Time-dependent (0 to 60 min) limited proteolysis (trypsin) of CHP3 (top) and the complex of CHP3:CBD (bottom) in the presence of both Mg^2+^ and Ca^2+^. Positions of full-length protein (FL) and two major proteolytic fragments (1 and 2) as well as CBD are indicated on the right of the Coomassie-stained SDS-PAGE gel, positions of co-separated molecular mass standards (mass in kDa) – on the left; the sample containing only trypsin was loaded on the first lane (T). (**B**) time-dependent limited proteolysis of CHP3 and mCHP3 in the presence of Mg^2+^, Ca^2+^, both ions or in the absence of them. Sections of the gel with bands corresponding to the full-length protein (FL) and two major proteolytic fragments (1 and 2) are shown. Nearly no degradation was observed for CHP3 and mCHP3 in the complex with CBD in all conditions (Mg^2+^+Ca^2+^ condition is presented here, other gels are shown in Figure S5A); (**C**) Schematic representation of full-length CHP3 with indication of N- and C-lobes, four EF-hand motifs (active EF-3 is highlighted in green). Proteolytic fragments 1 and 2 and trypsin cleavage sites were identified by mass spectrometry. (**D**) Ribbon presentation of the CHP3 AlphaFold2.0 model (40) with N- and C-lobes shown in blue and orange, respectively, and the connecting CHP-loop in cyan; the trypsin cleavage sites are highlighted in yellow and Ca^2+^ ion as a green sphere. The Ca^2+^ position in EF-3 was modelled by superimposition of the CHP3 model with the CHP1 X-ray structure, pdb ID 2ct9 (34).

We mapped both major cleavage sites on the 3D structure of CHP3 predicted with AlphaFold2.0 (40) (Figure 4D). This model closely resembles the structures of CHP1 and CHP2 in complex with CBD (42, 43), of CHP1 in complex with full-length NHE1 (44) and of CHP1 with an artificial C-terminal helix bound in the target-binding pocket (45), i.e. structures of target-bound CHPs. All these structures represent the Ca^2+^-bound state. The cleavage site R155/S156 is located in this model at the N-side of the incoming α-helix of EF-4 and though surface exposed the α-helical location may hamper the proteolytic digest (46). The other site, R73/K74 is located in the middle of the EF-2 loop (non-functional Ca^2+^ binding loop) (Figure 4D). Though the site appears to be surface exposed in the model, one should note that CHP3 has a 9-amino acid long insertion in that position as compared to CHP1 and CHP2 (Figure S6). The structure of CHP3 may deviate from the model and information for other conformations is lacking. Clearly, proteolysis showed that both sites are accessible for cleavage in the free CHP3 and become inaccessible upon complex formation. This proves substantial structural rearrangements upon target peptide binding that lead to overall stabilization of the protein.

### Ca^2+^ binding and the N-terminal myristoylation independently modulate the affinity of CHP3 for NHE1

To analyse how these conformational dynamics affect CHP3 function, we measured affinities of CHP3 and mCHP3 for the target peptide CBD with microscale thermophoresis (MST) in Ca^2+^- and in Mg^2+^-bound states. We used the system established earlier for quantification of binding affinities of CHP isoforms (CHP1, CHP2, CHP3) to NHE1, in which the target peptide CBD was fused to maltose-binding protein (MBP-CBD) for its stabilization (15, 33). We conducted the MST experiments with three biological replicates per sample and five individual titration series (technical replicates) per biological replicate (Figure 5). The dissociation constant (*K*_D_) for a given sample was calculated with a global parameter estimation procedure including all biological and technical replicates. We fitted the mass action law to the MST data points using non-linear regression. Confidence intervals were assessed for each *K*_D_ value by calculating profile likelihoods (error-surface plots) as described elsewhere (47, 48) (Figure 5E). Comparison of the resulting *K*_D_ values showed that the effects of Ca^2+^ binding and myristoylation on CHP3:CBD interaction are independent. Namely, Ca^2+^ binding increased affinities for CBD of both non-myristoylated (CHP3:CBD, *K*_D_=3.1 [2.8; 3.4] nM) and myristoylated CHP3 (mCHP3:CBD, *K*_D_=5.6 [5.1; 6.2] nM) by about fivefold as compared to the Mg^2+^-bound state (CHP3:CBD, *K*_D_=14.6 [12.5; 16.9] nM; mCHP3:CBD, *K*_D_=28.2 [26.6; 29.8] nM) (Figure 5A-D). In turn, for both Ca^2+^- and Mg^2+^-bound states, myristoylation decreased the affinity twofold (Figure 5). Hence, CHP3 binds to NHE1 with nanomolar affinity independently of the conformational state and modification, however, Ca^2+^ binding and myristoylation modulates the affinity.

**Figure 5.**
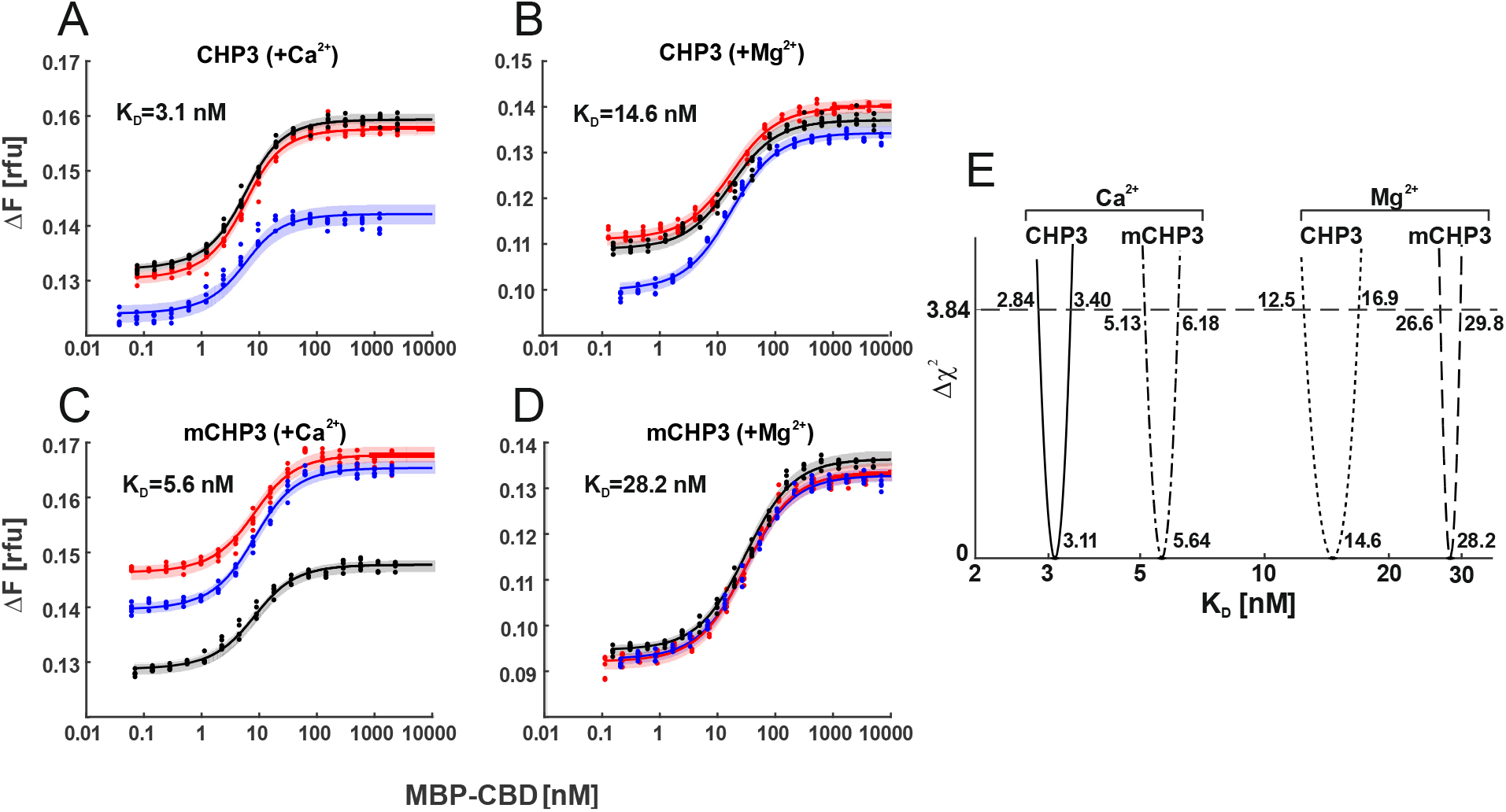
Ca^2+^ binding and myristoylation independently affect the interaction between CHP3 and MBP-CBD as measured with microscale thermophoresis. The interaction of CHP3 (**A-B**) and mCHP3 (**C-D**) with MBP-CBD was measured with three biological replicates shown in different colours (each with five technical replicates) in the presence of either Ca^2+^ (**A, C**) or Mg^2+^ (**B, D**). The combined data from individual experiments were fitted with a one-site binding model using global non-linear regression. (**E**) 95% confidence intervals (Δχ^2^ of 3.84) of *K*_D_’s for CHP3:CBD and mCHP3:CBD calculated with profile likelihood method. Raw MST traces are shown in Figure S7.

### Target peptide and Ca^2+^ binding regulate the interaction of CHP3 with lipid membranes

As we showed that the effects of Ca^2+^ and myristoylation on conformational changes of CHP3 and mCHP3 are independent, the hypothesis of a Ca^2+^-myristoyl switch for CHP3 (4, 23) had to be questioned. Therefore, we compared the state-dependent binding of CHP3 and of the Ca^2+^-myristoyl switch control protein recoverin to POPC:POPS (3:1 molar ratio) liposomes with a co-sedimentation assay. In line with previous studies (49), non-myristoylated recoverin (Rec) only weakly interacted with liposomes in both Mg^2+^- and Ca^2+^-bound states, whereas the interaction of myristoylated protein (mRec) increased about 4-fold upon Ca^2+^ addition (Figure 6A-B). In the closed Mg^2+^-bound state, CHP3 interaction with liposomes was similar as for recoverin, only a small protein amount co-sedimented with liposomes. In contrast to recoverin, Ca^2+^ addition increased the binding of non-myristoylated CHP3 (about 2.5 times), whereas myristoylation did not further increase the association of CHP3 with liposomes (Figure 6A-B). Interestingly, the binding of the target peptide CBD substantially changed the interaction of CHP3 with liposomes. In both the absence and the presence of Ca^2+^, only a small amount of the CHP3:CBD complex was associated with liposomes. However, CHP3 myristoylation increased the binding in both conditions about fourfold (Figure 6A-B). In the presence of Ca^2+^, the binding of mCHP3:CBD was even slightly higher than without it, however the difference was not significant. These results indicate the following mode of interaction. The myristic moiety is located in the protein core of target-free CHP3 and is not accessible to lipid membranes in the presence of Ca^2+^. Most likely, other protein regions are involved in membrane binding in this case. Target peptide binding causes conformational rearrangements with displacement of the myristic moiety from the protein core to the environment, which leads in consequence to an increased association of CHP3 and its target to lipid membranes.

**Figure 6.**
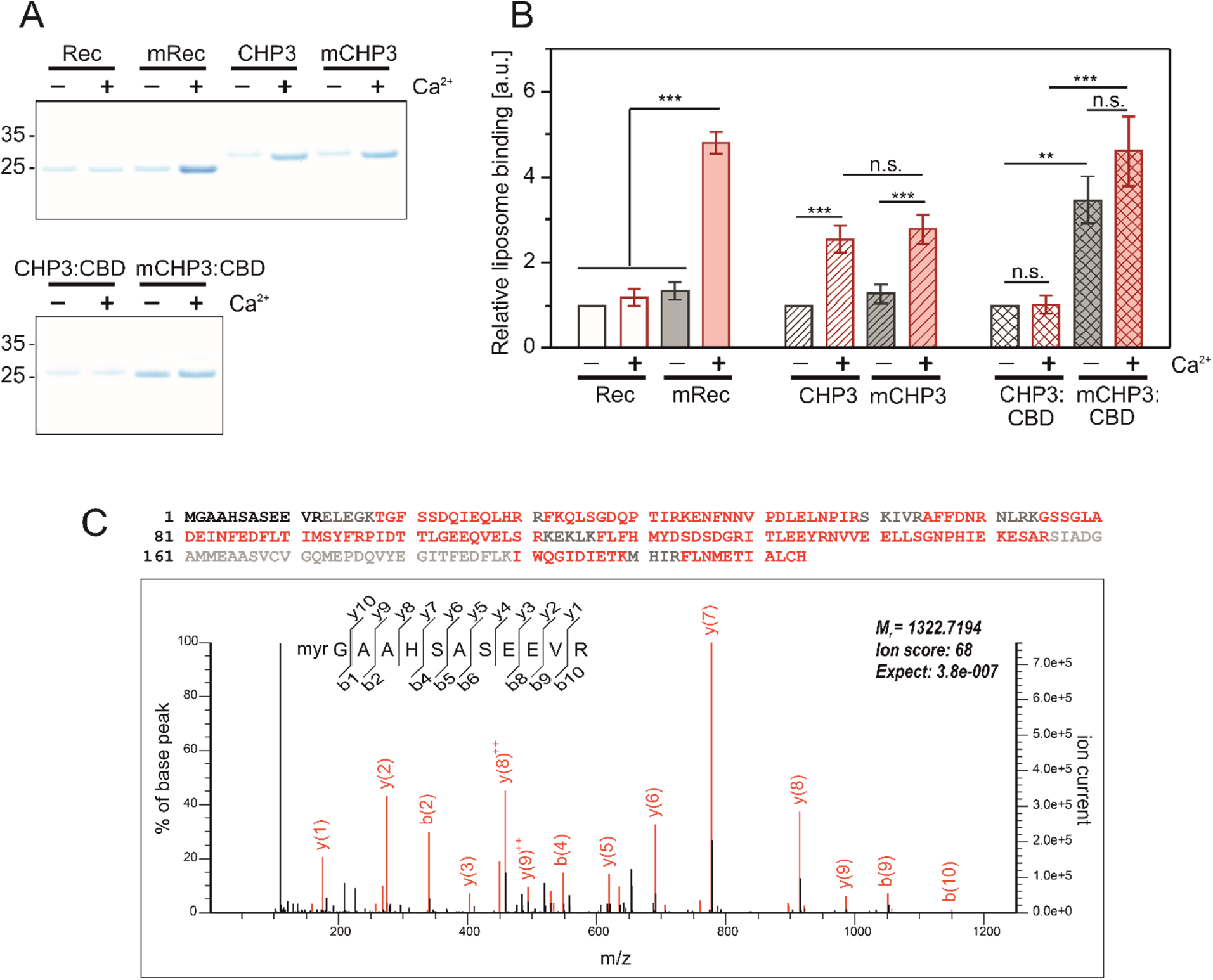
A-B) The interaction of CHP3 and mCHP3 with liposomes is regulated by Ca^2+^ and target peptide binding. Proteins were co-sedimented with POPC:POPS (3:1 molar ratio) liposomes in the presence of either 2 mM Mg^2+^ or 2 mM Mg^2+^ + 2 mM Ca^2+^ at 24°C. Non-myristoylated and myristoylated recoverin (Rec and mRec, respectively) were used as Ca^2+^-myristoyl switch control proteins. (**A**) Amount of proteins co-sedimented with liposomes was analysed with SDS-PAGE (4-12%, Bis-Tris). (**B**) Quantification of protein-liposome binding based on the densitometry of bands (SDS-PAGEs shown in **A**) corresponded of the co-sedimented proteins with three biological (CHP3 and complexes) or technical (recoverin) replicates. Values were normalized to the respective non-myristoylated protein in the Mg^2+^-bound state. Data are shown as mean±SD, one-way ANOVA with Tukey post-test was performed for mean comparison (statistical significance: n.s. – p>0.05, **p<0.01, ***p<0.001). **C) N-terminal myristoylation of target-associated CHP3 in mouse brain**. LC-MS/MS analysis of a trypsin-digested size fraction of solubilized mouse brain membrane containing NHE1-associated CHP3 (see Material & Methods). Upper panel: high coverage of the mouse CHP3 primary sequence by MS/MS-identified peptides (in red; black: sequences not identified, grey: sequences not accessible to our MS analysis) without inclusion of N-myristoyl modification. Lower panel: MS/MS spectrum from the same measurement assigned to the myristoylated tryptic N-terminal peptide of CHP3. Note that no other forms of the N-terminal peptide were detectable in error-tolerant search.

Finally, we investigated the N-terminal myristoylation status and membrane association of CHP3 *in vivo* using liquid-chromatography coupled mass spectrometry (LC-MS/MS). In three different preparations, namely (i) size fractionated complexes from mouse brain membranes (Fig. 6C), (ii) integral membrane proteins isolated from mouse platelets (i.e. after carbonate extraction, Figure S8), and (iii) anti-NHE1-affinity purification from solubilized mouse brain membranes (data not shown) we identified the vast majority of MS-accessible CHP3 peptides, but only the myristoylated form of the N-terminal peptide. Together, this suggests that the major form of CHP3 is both NHE1-associated and membrane-anchored in agreement with a target-dependent exposure and membrane integration of the N-terminal myristoyl moiety.

## Discussion

Ca^2+^-induced conformational changes are a hallmark of Ca^2+^-sensing EFCaBPs, which control the interaction with and function of specific target proteins (50). Based on the reported affinities of CHP3 for Ca^2+^ and Mg^2+^ (4), CHP3 should be able to react to changes of the intracellular calcium ion level with conformational changes. CHP3 binds Ca^2+^ with a *K*_D_ of 0.8 µM in the absence of Mg^2+^ and of 3.5 µM in the presence of 1 mM Mg^2+^ as determined by intrinsic tryptophan fluorescence (4). This distinguishes CHP3 from CHP1 and CHP2 isoforms that have much higher affinities for Ca^2+^ (*K*_D_ below 100 nM) (16, 17). In the absence of Ca^2+^, CHP3 binds Mg^2+^ with a *K*_D_ of 72 µM (4). Thus, CHP3 should bind Mg^2+^ in the resting state of the cell when the Ca^2+^ level is about 0.1 µM (18) and free Mg^2+^ concentration is maintained in the range of 0.25-1 mM (51). Upon a rise of the intracellular Ca^2+^ level, which can reach a concentration above 1 µM (18), Ca^2+^ binding to CHP3 could induce conformational changes allowing it to react to respective Ca^2+^ signals, as initially indicated by changes of the intrinsic tryptophan fluorescence of murine His_6_-CHP3 upon Ca^2+^ addition (4). Additionally, CHP3 can be N-terminally myristoylated and was proposed to be regulated with a so-called Ca^2+^-myristoyl switch mechanism (4, 23), in which Ca^2+^ binding induces the exposure of the myristic moiety to the environment. In contrast to this hypothesis, we describe in the present study independent effects of Ca^2+^ binding and N-terminal myristoylation on CHP3 conformation and on the interaction with its target protein NHE1 as well as a target-binding controlled association to lipid membranes.

First, we optimized the protein production and purification to obtain pure myristoylated and non-myristoylated CHP3 devoid of affinity tags, a prerequisite for further quantitative biochemical and biophysical analysis. Ca^2+^-dependent hydrophobic interaction chromatography (HIC) has been used for purification of EFCaBPs (36, 37), and we readily adapted it for the purification of both CHP3s. Myristoylated CHP3 (mCHP3) was produced by co-expression with the human NMT1 and supplementation of the culture medium with myristic acid as described for the production of N-myristoylated Nef protein of HIV-1 (52). It is still a challenging task to obtain stoichiometrically myristoylated recombinant EFCaBPs (53). However, with the expression system generated, we obtained preparative amounts of fully myristoylated CHP3 as confirmed by native MS analysis. With native MS, we also showed that both CHP3 and mCHP3 bind a single Ca^2+^, and are thus functional EFCaBPs, in agreement with the results of equilibrium dialysis shown earlier (4). Using the highly purified proteins, we showed with FPH and intrinsic fluorescence assays reversible conformational changes in CHP3 between a more hydrophobic Ca^2+^-bound state (open) and a less hydrophobic Mg^2+^-bound state (closed) (Figure 7). Such conformational changes are typical for Ca^2+^-sensor EFCaBPs including calmodulin, recoverin, neuronal calcium sensor protein (NCS1) and others, and serve as a conformational switch for binding and/or regulation of a target protein (54). Many EFCaBPs also bind Mg^2+^ that stabilizes them in a closed conformation in the absence of Ca^2+^ (55, 56). With nanoDSF analysis we revealed such a stabilization effect on CHP3, as Mg^2+^ increased its thermal stability. In contrast, Ca^2+^ binding slightly reduced the thermal and the proteolytic stability pointing towards an increased flexibility of the open state. The lower stability derived probably from an increased flexibility in the N-lobe of the protein, as we show that Ca^2+^ addition accelerated proteolysis within EF-2 located in the N-lobe. The apo-state of CHP3 appeared to be a destabilized form, which could not be reversibly converted into functional protein by Ca^2+^ supplementation. Thus, we conclude that Mg^2+^ binding is important to stabilize the protein in the native state at resting Ca^2+^ level.

**Figure 7.**
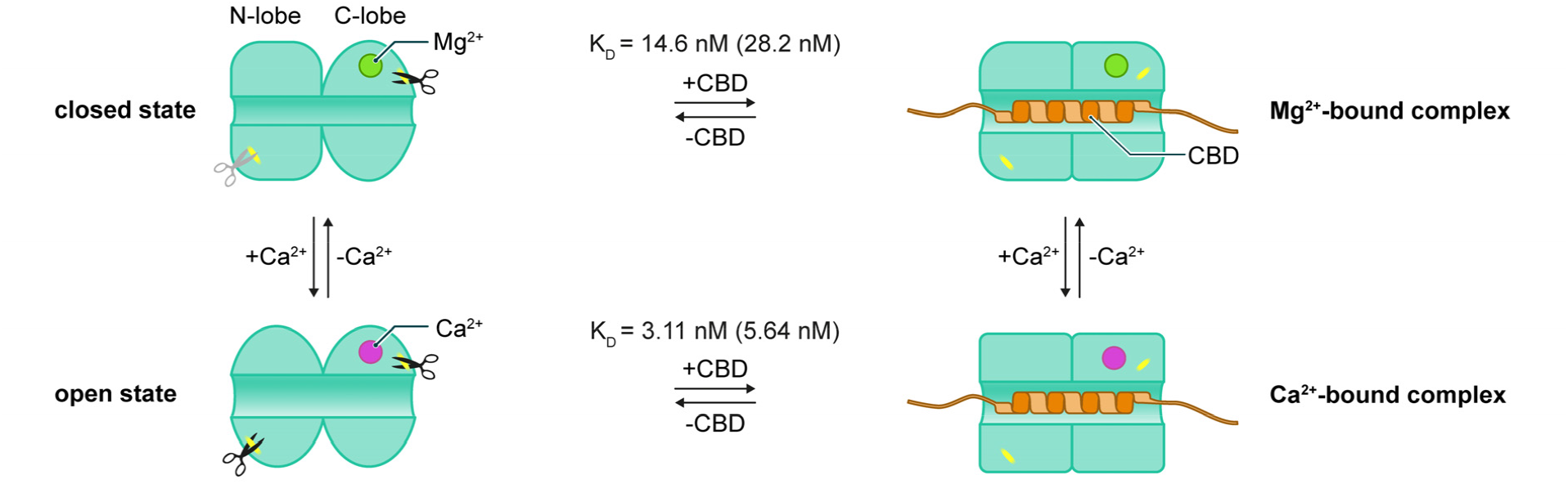
Conformational dynamics of CHP3 are controlled by Ca^2+^ and target peptide (NHE1 CBD) binding. Ca^2+^ (magenta) replaces Mg^2+^ (green) in EF-3 and induces the transition from the closed to the open state. Ca^2+^ binding increases the flexibility in the N-lobe accelerating the tryptic cleavage and reducing the thermal stability of CHP3. In both, Ca^2+^- and Mg^2+^-bound states, CHP3 binds the target peptide CBD with nanomolar affinity (K_D_ values indicated, in brackets for myristoylated CHP3). Complex formation has no effect on the thermal stability of the closed Mg^2+^-bound state, whereas it strongly enhances thermal stability of the open Ca^2+^-bound state. Independent of the bound ion, both trypsin cleavage sites are protected from proteolysis in the CHP3:CBD complex.

The Ca^2+^-induced higher flexibility of the open state appears to provide the molecular basis for CHP3 to interact more readily with its target(s) (Figure 7). Our data support this hypothesis. Firstly, the binding of the target peptide resulted in a drastic increase of CHP3 thermal and proteolytic stability. However, the thermal stability increased only in the presence of Ca^2+^, whereas for the Mg^2+^-bound state it remained unchanged suggesting a lower energy state of the target bound form in the presence of Ca^2+^. The higher flexibility of the open Ca^2+^-bound state may lower the energy barrier for binding of the target peptide. Secondly, in line with co-immunoprecipitation results for tagged NHE1 and tagged CHP3 from transfected AP-1 cells (23), we detected binding between CHP3 and CBD in both, Ca^2+^- and Mg^2+^-bound states (see Figure 5). The quantitative analysis with MST showed that the affinity of the open Ca^2+^-bound state was about five times higher, clearly indicating that Ca^2+^ controls CHP3 function.

Though all CHP isoforms bind NHE1 with nanomolar affinities in the absence and presence of Ca^2+^ (15, 33), intracellular Ca^2+^ signals might regulate NHE1 function (e.g. activity or stability) via CHPs affecting the conformation of the complex. Regulation of the activity by Ca^2+^ binding to the regulatory subunit was described for calcineurin, a phosphatase consisting of the catalytic and regulatory subunits calcineurin A (CnA) and B (CnB), respectively. Similar to the CHP3:NHE1 interaction, CnB forms a stable complex with CnA independently of Ca^2+^ binding. Nevertheless, Ca^2+^ binding induces conformational changes of CnB resulting in the release of the calmodulin-binding domain of CnA and allowing its activation by calmodulin (19, 57). In the case of CHPs, and particularly of CHP3, Ca^2+^ binding to the C-terminal EF-hands (EF-3 in CHP3) affects the conformation also of the N-terminal lobe (Figure 7), most likely via the hydrophobic cluster at the interface between N- and C-lobes, which is characteristic for the so-called CPV (calcineurin B, p22/CHP1, visinin) subfamily of EFCaBPs (58). Such conformational changes of CHP might affect the overall structure of the complex formed between CHPs and NHE1. All CHP structures so far, the recent cryo-EM structure of the NHE1:CHP1 complex (44), and structures of CHP1 (43) and CHP2 (42) in complex with the target peptide CBD obtained with NMR and X-ray crystallography, showed CHPs in the Ca^2+^- and target peptide bound state (or with the Ca^2+^-mimic ion Y^3+^ in the CHP2:CBD complex). The structure of CHP3 in any state has not been determined so far and moreover, structural information of CHPs in the Ca^2+^-free state (Mg^2+^-bound state) and in open, target-free conformation is lacking. Therefore, further structural studies are required to better understand the molecular mechanisms for NHE1 regulation by Ca^2+^ binding to CHPs.

Regarding the lipidation modification, many Ca^2+^ sensor proteins have an N-terminal myristoylation site and often Ca^2+^ binding to the myristoylated EFCaBP causes an exposure of the myristoyl moiety. A surface-exposed myristoyl moiety strongly increases the binding of the Ca^2+^-bound EFCaBP to lipid membranes. This regulatory mechanism is called Ca^2+^-myristoyl switch (59). A Ca^2+^-myristoyl switch was proposed for CHP3, as mutational analysis indicated that deficiency in Ca^2+^ binding and/or myristoylation of CHP3 reduced the surface expression and activity of NHE1 in the same degree (23). Yet, direct evidence for a Ca^2+^-myristoyl switch in CHP3 was lacking. Our data indicate an independent regulation of CHP3 by Ca^2+^ binding and myristoylation, and contradict the presence of a Ca^2+^-myristoyl switch. Firstly, CHP3 myristoylation only slightly increased the hydrophobicity of the Ca^2+^-bound state in comparison to the prominent hydrophobicity increase of recoverin that has a classical Ca^2+^-myristoyl switch (59). Secondly, myristoylation did not affect Ca^2+^-binding properties of CHP3. Next, myristoylation reduced the thermal stability of free and target-bound CHP3 in a similar manner for both Ca^2+^- and Mg^2+^-bound states. Furthermore, mCHP3 affinities to CBD in both ion-bound states were decreased twofold compared to CHP3. We also provided further evidence for the lack of a Ca^2+^-myristoyl switch in CHP3 by analyzing its interaction with liposomes. Although Ca^2+^ addition strongly increased the CHP3 association with POPC/POPS liposomes, the effect was the same for both myristoylated and non-myristoylated proteins. In the closed conformational state, the myristic group is usually buried in the protein core of EFCaBPs, however the binding interface differs a lot in different EFCaBPs (22). We assume that CHP3 resembles guanylyl cyclase activating proteins (GCAPs), which have the myristoyl group enclosed in the protein hydrophobic core independently of Ca^2+^ binding (22). The membrane binding of CHP3 independent of its myristoylation suggests that different protein regions, e.g. clusters of hydrophobic or positively-charged residues, are involved in protein-lipid interaction. Binding of the target peptide CBD drastically reduced the interaction of CHP3 with lipid membranes apparently caused by large conformational rearrangements. The CHP3:CBD complex did not bind to liposomes in the presence of Ca^2+^. One can speculate, that residues involved in membrane binding of the target-free Ca^2+^-bound CHP3 become inaccessible upon complex formation. In stark contrast to the target-free CHP3, myristoylation increased fourfold the amount of CHP3:CBD complex associated with liposomes. We propose that the target peptide competes with the myristoyl moiety for binding in the hydrophobic pocket and by this way shifts the conformation equilibrium of CHP3, resulting in the exposure of the myristoyl moiety to the environment (Figure 8). The competition for the hydrophobic pocket is also in line with the lower affinity of mCHP3 to CBD in comparison to non-myristoylated protein as shown by MST analysis (Figure 5). In general, the accessibility of the myristoyl moiety in myristoylated regulatory proteins can be induced by Ca^2+^ or by small molecule binding, phosphorylation and pH shift. Respective regulatory mechanisms were termed Ca^2+^-myristoyl switch, ligand-dependent, phosphorylation (24), and pH-dependent (29) switches. The myristoyl switch caused by protein-protein interaction described here for CHP3 is a novel mechanism that regulates the association of myristoylated proteins with lipid membranes. We propose to call it “target-dependent myristoyl switch” (Figure 8). This mechanism was confirmed by LC-MS/MS analyses of CHP3-associated complexes in brain (Figure 6C) and a preparation from platelets containing integral membrane proteins and lipid-anchored proteins (Figure S8) which exclusively detected the N-terminally myristoylated CHP3. We suggest that the target-dependent myristoyl switch is the canonical mechanism to anchor the regulatory domain of NHE1 to the plasma membrane, which appears to be the molecular basis for an increase of NHE1 surface stability by CHP3. A defined conformation enforced by the membrane anchor may also regulate NHE1 activity. As second mode of regulation, Ca^2+^ binding to CHP3 can affect NHE1:CHP3 interaction in two ways. Ca^2+^ directly increases the binding affinity of CHP3 for NHE1, and indirectly increases the probability to interact with NHE1 via enhanced association of free CHP3 with lipid membranes. Our hypothesis is in line with the previous observation that both, CHP3 myristoylation and Ca^2+^ binding are important for NHE1 surface stability when both proteins were co-expressed in AP-1 cell line (23).

**Figure 8.**
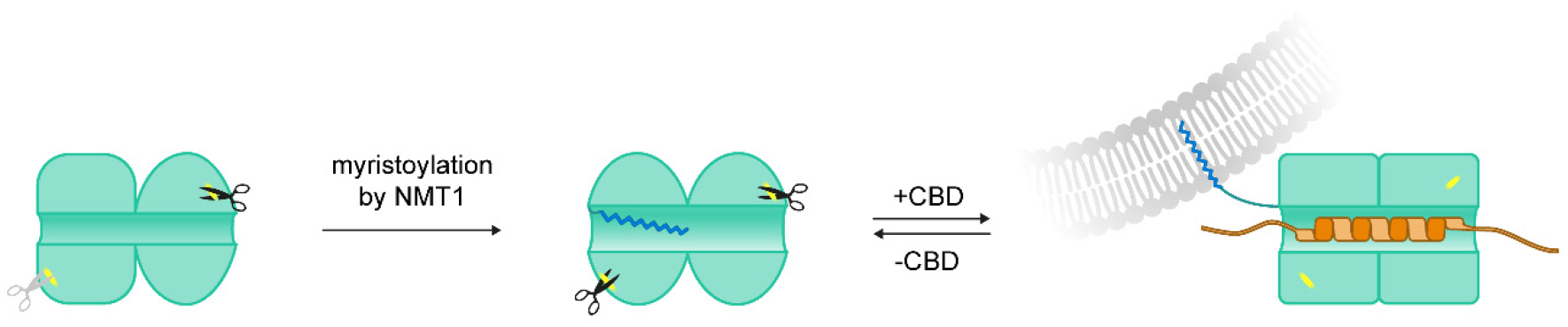
Target-dependent myristoyl switch in CHP3. N-terminal myristoylation of CHP3 by NMT1 increases the flexibility in the N-lobe, as it accelerated its tryptic cleavage. The myristoyl moiety most likely partially occupies the target binding pocket in both Ca^2+^- and Mg^2+^-bound states, as myristoylation reduced the affinity of CHP3 to CBD. CBD binding causes displacement of the myristoyl moiety from the hydrophobic pocket, as the surface exposed myristoyl moiety enhanced CHP3 binding to lipid membranes in both Ca^2+^- and Mg^2+^-bound states.

It will be interesting to investigate whether the interaction of CHP3 with other described target proteins, such as with subunit 4 of COP9 signalosome, GSK3ß and calcineurin A (4-6) can induce the target-dependent myristoyl switch and how Ca^2+^ binding affects these interactions. Structural characterization of CHP3 in different conformational states is needed to further explore the underlying molecular basis of the regulatory mechanism. The target-dependent myristoyl switch might exist in other proteins including CHP1 and CHP2 and can be explored by comparison of the membrane association for myristoylated and non-myristoylated complexes.

CHP3 is an emerging important player in the cellular Ca^2+^ signalling network that is involved in regulation of cancerogenesis, cardiac hypertrophy and neuronal development. Our data can help to better understand the molecular mechanisms of regulation of these processes by CHP3.

## Material and Methods

### Plasmids

The generation of all plasmids was performed using NEBuilder HiFi DNA assembly Master Mix (New England Biolabs GmbH, Frankfurt, Germany). Protein coding sequences (CDS) were inserted without any additions for affinity tags if not stated otherwise. The pETDuet-1_NMT1_Nef plasmid containing the codon optimized CDS of the human N-myristoyl transferase 1 (NMT1, Gene ID 4836, UniProtKB ID P30419) was kindly provided by Prof. Willbold (Forschungszentrum Jülich, Germany). To obtain the pETDuet_NMT1_TESC plasmid, the codon optimized (*E. coli*) CDS of human CHP3 (TESC, Gene ID 54997, UniProtKB ID Q96BS2) was cloned into the second multiple cloning site of the pETDuet-1_NMT1_Nef plasmid replacing the Nef CDS. The NMT1 CDS was deleted from the pETDuet-1_NMT1_TESC plasmid to generate the plasmid pETDuet1_TESC. For co-expression of the NHE1 CBD:CHP3 complexes, the codon optimized (*E. coli*) CDS of a minimal NHE1 CBD with three additional histidines added C-terminally (NHE1 residues 515-545, MRSINEEIHTQFLDHLLTGIEDICGHYGHHHHHH, CBDHis) was used. In case of the pETDuet_CBDHis_TESC plasmid, the CBDHis CDS was inserted in the first multiple cloning site, whereas for the pETDuet_NMT1_CBDHis_TESC plasmid, the CBDHis CDS and the linker region containing T7 promoter/lac operator and ribosome binding site was inserted between NMT1 and CHP3 CDS’s. The plasmid pETDuet1_Rec containing the codon optimized CDS of human recoverin (Gene ID 5957, UniProtKB ID P35243) in the second multiple cloning site was obtained from BioCat (Heidelberg, Germany). NMT1 CDS was inserted into the first site of this plasmid to obtain the pETDuet1_NMT1_Rec plasmid.

### Protein productions and purifications

We adapted the protocol of Glück *et al*. (52) for the production of recombinant myristoylated CHP3 (mCHP3). Pre-cultures in TB^carb^ medium were made from a single colony of E. *coli* BL21 (DE3) transformed with the pETDuet_NMT1_TESC plasmid. Main cultures in TB^carb^ medium containing the surfactant Antifoam 204 (Merck KGaA, Darmstadt, Germany) were inoculated to a final OD_600_=0.1 and incubated at 37°C. At OD_600_=0.6, the temperature was shifted to 28°C and 3% ethanol (v/v) was added. Myristic acid was added at OD_600_=0.70-0.75. The protein production was induced at OD_600_=0.8 with 0.5 mM IPTG. After 4 h incubation, cells were harvested and stored at -80°C. Non-myristoylated CHP3 was produced similarly using the plasmid without NMT1 (pETDuet_TESC); no antifoam, ethanol and myristic acid were added to the medium. The expression was performed at 30°C overnight.

All chromatography columns were obtained from Cytiva (Marlborough, MS, USA). CHP3 and mCHP3 were purified using Ca^2+^-dependent hydrophobic interaction chromatography based on protocols established for other EFCaBPs (36, 37). Briefly, sedimented cells were resuspended in buffer A (20 mM HEPES, 150 mM NaCl, 10 mM DTT, 10 % glycerol (w/v), pH 7.2) containing 10 mM CaCl_2_ and 0.3 mM Pefabloc (Carl Roth GmbH, Karlsruhe, Germany), and lysed with a cell disruptor (Constant systems, Daventry, UK). Cell debris was removed by centrifugation and the supernatant was loaded onto a HiPrep 16/10 Phenyl HP column equilibrated with buffer A containing 10 mM CaCl_2_. After removal of unbound material with 10 column volumes of the equilibration buffer, the protein was eluted with buffer A containing 10 mM EGTA, and the eluate was mixed with an equal volume of buffer A containing 10 mM MgCl_2_. The eluted protein was concentrated and loaded onto a pre-equilibrated HiLoad 26/600 Superdex 75 column. Buffer A containing 10 mM MgCl_2_ was used as running buffer. Fractions containing the purified protein were pooled, concentrated, mixed 1:1 with the cryo-storage buffer (2M sucrose in gel filtration buffer), flash frozen in liquid nitrogen, and stored at -80°C. Recoverin and mRecoverin were produced and purified as described above for CHP3 and mCHP3 using the pETDuet_Rec and pETDuet_NMT1_Rec plasmids, respectively. The fusion construct of maltose binding protein and NHE1 CBD (MBP-CBD) was produced and purified as described previously (33).

The production of CHP3:CBD and mCHP3:CBD complexes was performed as described above using pETDuet_CBDHis_TESC or pETDuet_NMT1_CBDHis_TESC plasmids, respectively. Cells were lysed in the presence of 20 mM imidazol. The complexes were purified by IMAC followed by gel filtration. Cell lysate was loaded onto a HisTrap FF 5 ml column pre-equilibrated with buffer A containing 10 mM CaCl_2_ and 20 mM imidazol. After removal of unbound material with 10 column volumes of the equilibration buffer, elution was done with a linear gradient from 20 to 600 mM imidazol over 20 column volumes. For gel filtration, buffer A containing 10 mM CaCl_2_ was used as running buffer.

### Native mass spectrometry

The degree of myristoylation was determined by native MS. For native ion mobility separation (IM)-MS analysis, proteins were transferred into 200 mM ammonium acetate, pH 6.7. Capillaries for nanoflow electrospray ionization were prepared in-house from borosilicate glass capillaries 1.0 mm OD x 0.78 mm ID (Harvard Apparatus, Holliston, MA, USA) using a micropipette puller Model P-97 (Sutter Instruments, Novato, CA, USA) and coated with gold using a sputter coater 108 auto (Cressington TESCAN GmbH, Dortmund, Germany). 3-5 µl of 20 µM protein were loaded per experiment. Measurements were carried out with quadrupole-IM-MS-ToF instrument Synapt G1 HDMS (Waters Corporation, Milford, MA, USA and MS Vision, Almere, The Netherlands) equipped with a 32,000 *m/z* range quadrupole. Pressures were 8 mbar backing in the source, 0.017 mbar argon in the trap and transfer collision cell and 0.5 mbar nitrogen in the ion mobility cell. Instrument parameters were as follow: capillary voltage 1.2 kV, source temperature 60°C, sample cone voltage 100 V, extraction cone voltage 10 V, acceleration voltages in the trap and transfer cell 5 V and 35 V, respectively, IM bias potential 22 V, wave velocity and height 300 m/s and 8 V. Spectra were acquired in the positive ion mode in the *m/z* range 1000-6000. Spectra recorded for cesium iodide at 50 mg/ml in 50% (v/v) isopropanol were used for external mass calibration. Data were processed and analysed using MassLynx Software version 4.1 (Waters Corporation) and UniDec version 2.7.3 (60). For charge state deconvolution, smoothing and subtraction of linear background from raw spectra were done.

### MS sequencing of native CHP3

Mouse brain membranes were prepared, solubilized with ComplexioLyte 47 (Logopharm, Germany) and protein complexes were separated by BN-PAGE as described (61). A section corresponding to an apparent molecular weight of 630 kDa (the migration size of native NHE1 complexes) was excised and subjected to MS analysis (as described below). Mouse platelets were prepared from fresh blood by differential centrifugation and 20 µg washed with 50 µl 100 mM Na_2_CO_3_ pH 11 (10 min at 4°C, followed by ultracentrifugation at 233,000 x g for 20 min). The resulting pellet was dissolved in Laemmli buffer and separated on an SDS-PAGE gel and silver-stained. The section <50 kDa was excised, in-gel digested with trypsin, the peptides were dissolved in 20 µl 0.5% (v/v) trifluoroacetic acid, and 1 µl was analyzed on a LC-MS/MS setup (UltiMate 3000 RSLCnano HPLC / QExactive HF-X mass spectrometer, both Thermo Scientific, Germany) as described (62). MS/MS data was extracted and searched against the UniProtKB/Swiss-Prot database (mouse, rat, human, release 20220525; Mascot search engine version 2.7, Matrix Science, UK) with the following settings: Acetyl (Protein N-term), Carbamidomethyl (C), Gln->pyro-Glu (N-term Q), Glu->pyro-Glu (N-term E), Oxidation (M), Propionamide (C) as variable modifications, one missed cleavage allowed, with or without myristoylation (N-term G) or in error tolerant mode; precursor / fragment mass tolerance was ±10 ppm / ±25 mmu), significance threshold p<0.05.

### Fluorescence-based assays

The conformational changes of the protein in response to ion binding were probed with the fluorescence probe hydrophobicity (FPH) assay (15) and by monitoring the intrinsic tryptophan fluorescence. The buffer of the protein solution was exchanged to the assay buffer (20 mM HEPES, 150 mM NaCl, 1 mM TCEP, 2 mM MgCl_2_, pH 7.2) containing 1 mM EGTA using Zeba™ spin desalting columns (Thermo Fisher Scientific Inc., Waltham, MA, USA). The FPH assay was modified as follows. 500 µL of 1.5 µM protein and 5 µM ProteOrange (Lumiprobe GmbH, Hannover, Germany) in the assay buffer were mixed and incubated for 30 min in the dark. The fluorescence at 585 nm (excitation at 470 nm) was measured continuously with the Cary Eclipse spectrofluorimeter (Agilent Technologies, Santa Clara, CA, USA). Every 60 s a new compound was added in the following order: 2 mM CaCl_2_, 3 mM EGTA, 3 mM EDTA, and 4 mM CaCl_2_.

Determination of Ca^2+^ EC50 values was performed using the FPH assay, mixing 1.5 µM protein and 5 µM ProteOrange with varied (0.34 µM to 8.0 mM) concentrations of CaCl_2_. The assay was carried out in 96 well black plates (Corning, Corning, NY, USA) with a sample volume of 100 μl. For each biological replicate, three to four technical replicates were measured. The fluorescence at 590 nm with excitation at 485 nm (F_590_) was measured with the BioTek Synergy 2 plate reader (Agilent Technologies, Santa Clara, CA, USA). The data were fitted using the Hill equation

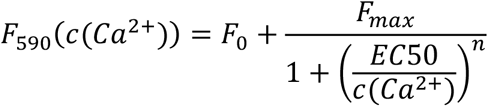

to estimate parameters from data which has the *a priori* unknown parameters: *F*_*0*_ (F_590_ in the absence of Ca^2+^), *Fmax* (maximal change of F_590_), *EC50* and *n* (Hill coefficient). The parameters *EC50* and the Hill coefficient *n* represent biological quantities that are global and therefore shared between the three biological replicates. The parameters *F*_*0*_ and *Fmax* are affected by experimental procedures and therefore different between the three biological replicates. Adding one parameter for dose-independent measurement noise, a total of nine parameters has to be estimated simultaneously for each condition. Parameter uncertainty can be assessed using the profile likelihood method, which is also referred to as error surface plots. We utilized optimization to obtain the best parameter set that minimizes deviation between model and data in terms of the negative log-likelihood. Uncertainty of the parameters are quantified using profile likelihood method (47), which is also termed error surface plots in the literature. This fitting procedure was independently applied to both conditions in order to obtain EC50 values and corresponding uncertainties for both conditions. Data2Dynamics Software (63) used for EC50 determination from data including biological and technical replicates is available at GitHub (https://github.com/Data2Dynamics/d2d).

For monitoring the intrinsic tryptophan fluorescence, 500 µL of 2.5 µM of the respective protein in assay buffer were prepared. Fluorescence was measured at 330 nm with an excitation at 280 nm. The time traces were recorded as described above for the FPH assay.

All assays were performed with three individually purified biological replicates of CHP3 and mCHP3.

### Thermal stability analysis with nanoDSF

The thermal stability of CHP3, mCHP3, and the complexes with CBD were compared at different conditions using nano differential scanning fluorimetry (nanoDSF). Buffer of the protein solutions were exchanged to the assay buffer as above; the final concentration was adjusted to 100 µM and thermal stability was measured in the presence of 10 mM of either MgCl_2_, CaCl_2_, MgCl_2_ and CaCl_2_ or EDTA. Each sample was measured with two technical and three independent biological replicates in standard capillaries (Nanotemper Technologies, Munich, Germany) with the Prometheus device (Nanotemper Technologies). Measurements were carried out from 20°C to 95°C with a heating rate of 1°C/min.

### Limited proteolysis

Next, we probed the proteolytic stability of CHP3, mCHP3 and of their complexes with CBD. Buffer of the protein solutions was exchanged to the assay buffer containing either 2 mM MgCl_2_+ 0.5 mM EGTA, 2 mM CaCl_2_, 2 mM MgCl_2_ + 2 mM CaCl_2_ or 2 mM EDTA as an additive. The final protein concentration was adjusted to 0.5 mg/ml for each sample. Each reaction was started with the addition of 0.01 mg/ml trypsin (Sigma Aldrich, Cat.No. T1426) dissolved in 1 mM HCl, and performed for 1 hour on ice. Samples were taken after 5, 15, 30, 45, and 60 min, directly mixed with an equal volume of 2x reducing NuPAGE™ LDS sample buffer (Thermo Fisher Scientific) and boiled for 5 min at 95°C. The proteolytic fragments were separated on 12 % NuPAGE Bis-Tris gels (Thermo Fisher Scientific).

To determine which trypsin cleavage sites in CHP3 are accessible for proteolysis, the major proteolytic fragments were analysed by ESI mass spectrometry. 100 µL of CHP3 and mCHP3 in the assay buffer with 2 mM CaCl_2_ and 2 mM MgCl_2_ were incubated with trypsin for 15 min. Protein fragments were separated on 16% Novex tricine gels (Thermo Fisher Scientific). The major bands were cut, minced into smaller pieces and loaded on top of 300 kDa Nanosep centrifugal devices (Pall Corporation, Port Washington, NY, USA). Proteins were eluted two times with 400 µL of the assay buffer containing 0.1 % SDS by centrifugation at 14,000g for 20 min. Eluates were concentrated with Vivaspin500 5,000 MWCO concentrators (Sartorius AG, Goettingen, Germany). SDS was removed by precipitation with 0.2 M KCl followed by 10 min centrifugation at 14,000g. 10 µL of each sample were analysed with liquid chromatography (Thermo Fisher Ultimate 3000 with Phenomenex Jupiter 5 µm C4 300 Å 50 × 2 mm) coupled to high resolution electrospray ionization (ESI) mass spectrometry (Maxis 4G, Bruker Corporation, Billerica, MA, USA). The resulting peptide masses were compared to all possible trypsin digestion fragments of CHP3 or mCHP3 to identify the cleavage sites.

### Microscale thermophoresis

Microscale thermophoresis was performed using the Monolith Pico device (Nanotemper Technologies) as described previously (33). The only difference was the use of the measuring buffer containing either 10 mM CaCl_2_ or 10 mM MgCl_2_ (20 mM HEPES, 5 mM TCEP, 150 mM NaCl, 0.05 % Tween-20, 10 mM MgCl_2_ or 10 mM CaCl_2_; pH 7.2). All measurements were performed with three independently produced and purified protein replicates of CHP3, mCHP3 and MBP-CBD. Every condition was titrated and measured five times per biological replicate (technical replicates).

Mass action kinetics of a one-site binding model was utilized to derive the fitting function for dissociation constant (K_D_) determination: Let A and B* denote the concentrations of unlabelled and labelled binding partners respectively, and the concentration of the protein complex by AB*. The total amounts of A and B* (*A*_*tot*_ and 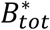) can be controlled by the experimental setup. In our application, 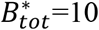 nM was fixed and *A*_*tot*_=[MBP-CBD] was the independent variable for the dose response analysis. The thermophoresis data were normalized and fitted with global non-linear regression to the dose response function (48):

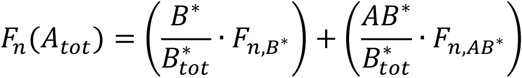

with *F*_*n*_(*A*_*tot*_) – thermophoresis (change of normalized fluorescence) as a function of MBP-CBD concentration, *F*_*n,B*∗_ and *F*_*n,AB*∗_ – thermophoresis of the labelled partner and the complex, respectively. The concentration of the complex is given by

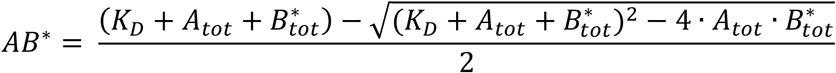

which follows from solving a quadratic function originating from mass action kinetics combined with equilibrium assumption for the signal strength.

Simultaneously, all individual parameters (*B*^∗^, *F*_*n,B*∗_, *F*_*n,AB*∗_) per biological replicate were fitted together with the global *K*_D_ value. To estimate the noise level within the technical replicates we used an error model with absolute and relative errors represented by two parameters per biological replicate. With three biological replicates, this results in a total of 16 parameters that were fitted simultaneously. We utilized deterministic multistart optimization to obtain the best parameter set that minimizes deviation between model and data in terms of the negative log-likelihood. Uncertainty of the parameter of interest *K*_D_ is quantified using profile likelihood method (47), which is also termed error surface plots in the literature. This fitting procedure was independently applied to each condition in order to obtain *K*_D_ values and corresponding uncertainties for all four conditions. Data2Dynamics Software (63) used for *K*_D_ determination from data including biological and technical replicates is available at GitHub (https://github.com/Data2Dynamics/d2d).

### Interaction of proteins with lipid membranes

The POPC:POPS (Avanti Polar Lipids, Birmingham, AL, USA) mixture (3:1 molar ratio) in chloroform was dried in round bottom glass vials under a nitrogen stream. The dried lipid film was resuspended in liposome buffer (20 mM HEPES, 150 mM NaCl, 10 % sucrose, pH 7.2) at a final concentration of 4 mM, sonified 3 times for 20 s with 40 s breaks (Bandelin Sonorex ultrasonic bath). Then liposomes were subjected to 10 freeze-thaw cycles and extruded through a 1 μm polypropylene membrane (Avestin, Ottawa, Canada). The size of the resulting liposomes was controlled with DLS (DynaPro NanoStar, Wyatt Technology, Santa Barbara, CA, USA). Liposomes were stored at room temperature in the dark.

The lipid-protein co-sedimentation assay was performed mostly as described previously (49). Briefly, proteins were transferred to the assay buffer. Liposomes were washed once with the assay buffer containing either 2 mM CaCl_2_ (Ca^2+^ condition) or 1 mM EGTA (Mg^2+^ condition) and mixed with the protein in the respective buffer. Final lipid and protein concentrations were 1.4 mM and 10 µM, respectively. The samples were incubated for 20 min at 24°C with constant mixing and centrifuged at 17,000g for 15 min. The pellet was washed once with the respective buffer and resuspended in 1x reducing NuPAGE LDS sample buffer (Thermo Fisher Scientific). Samples were analysed using 4-12% NuPAGE Bis-Tris gels (Thermo Fisher Scientific).

The lipid-binding experiments were performed with three individually purified biological replicates of CHP3 and mCHP3 and respective complexes with CBD. Three technical replicates were analysed for Recoverin and mRecoverin as controls. Densitometry of protein bands was done with ImageJ (NIH). Band intensities were normalized to the Mg^2+^ condition of the respective non-myristoylated protein.

### Data availability

All data generated or analyzed during this study are included in the manuscript. Data2Dynamics Software used for EC50 and K_D_ determination is available at GitHub (https://github.com/Data2Dynamics/d2d).

## Acknowledgements

This work was supported by the Deutsche Forschungsgemeinschaft (DFG, German Research Foundation) under Germany’s Excellence Strategy (CIBSS – EXC-2189 – Project ID 390939984) in the form of project funding to B.W, B.F., C.H., C.K. and of CIBSS Launchpad Program Funds to E.M.; by the DFG through Project-ID 403222702 / SFB 1381 to C.H., U.S. and B.W., through Project-ID 431984000 / SFB 1453 to B.F. and C.H., and through project-ID 278002225 / RTG 2202 to C.H.; and by the German Ministry of Education and Research by grant EA:Sys (FKZ031L0080) to C.K.. We thank Prof. Willbold (Forschungszentrum Jülich, Germany) for kindly providing the pETDuet-1_NMT1_Nef plasmid, Dr. Michael Pfeffer (MS Service at University of Basel) for performing ESI mass spectrometry, and Michal Rössler (CIBSS, University of Freiburg) for supporting the artwork in Figures 7-8.

## Author Contributions

FB, SF, EVM, CH conceived the study; FB and SF prepared the proteins; FB performed SDS-PAGEs, fluorescence assays, limited trypsinolysis and protein-liposome binding; SF performed MST; FB and SF performed nanoDSF; SL and EVM contributed to the development of the FPH assay; SF, JIH and SCH contributed to measurements of protein-liposome binding; LR and CK performed statistical analysis of MST and Ca^2+^ EC50 data; FD and BW performed native mass spectrometry and analyzed the data; WB, US and BF performed LC-MS/MS spectrometry and analyzed the data; FB, SF, EVM, CH analyzed all data; FB, SF, EVM, CH wrote the manuscript; all authors discussed the results and contributed to review and editing of the manuscript.

