## Supplemental Data for "Conformational dynamics and target-dependent myristoyl switch of calcineurin B homologous protein 3"

*contributed equally;

**Running title: Conformational dynamics of CHP3**

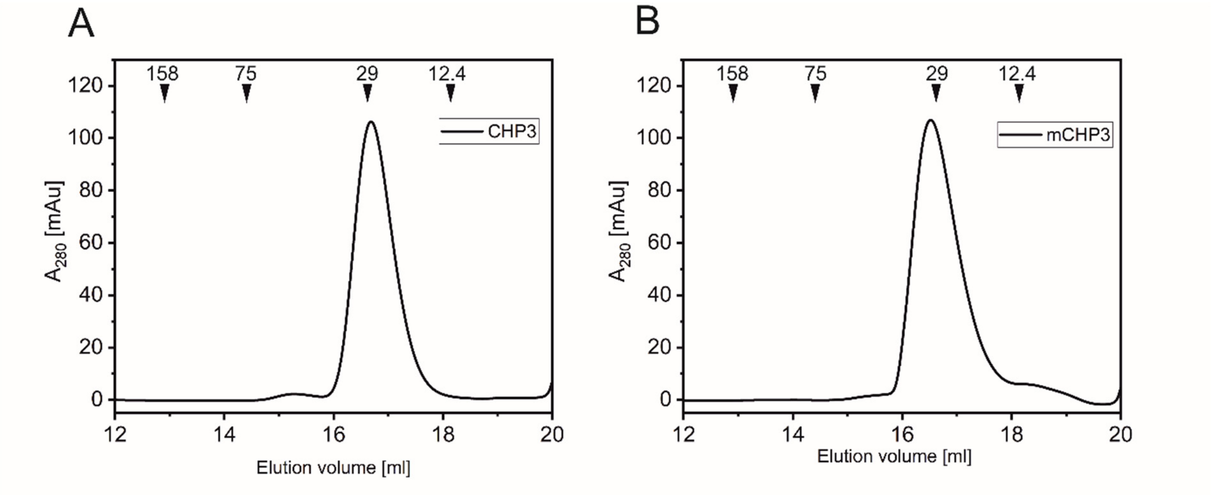


**Figure S1** Analytical size exclusion chromatography of purified CHP3 (**A**) and mCHP3 (**B**) on Superdex 75 Increase 10/300 GL. The position of elution peaks of mass standards (aldolase: 158 kDa, conalbumin: 75 kDa, carbonic anhydrase: 29 kDa, cytochrome c: 12.4 kDa) are indicated on the top.

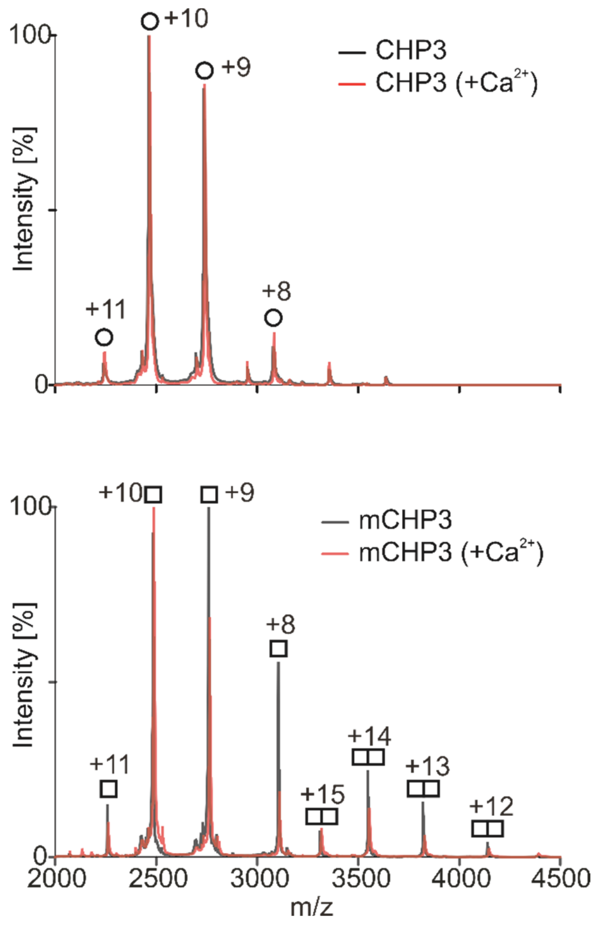


**Figure S2** Non-deconvoluted mass spectra showing charge state distributions of CHP3 and mCHP3 measured in the positive ion mode by native MS.

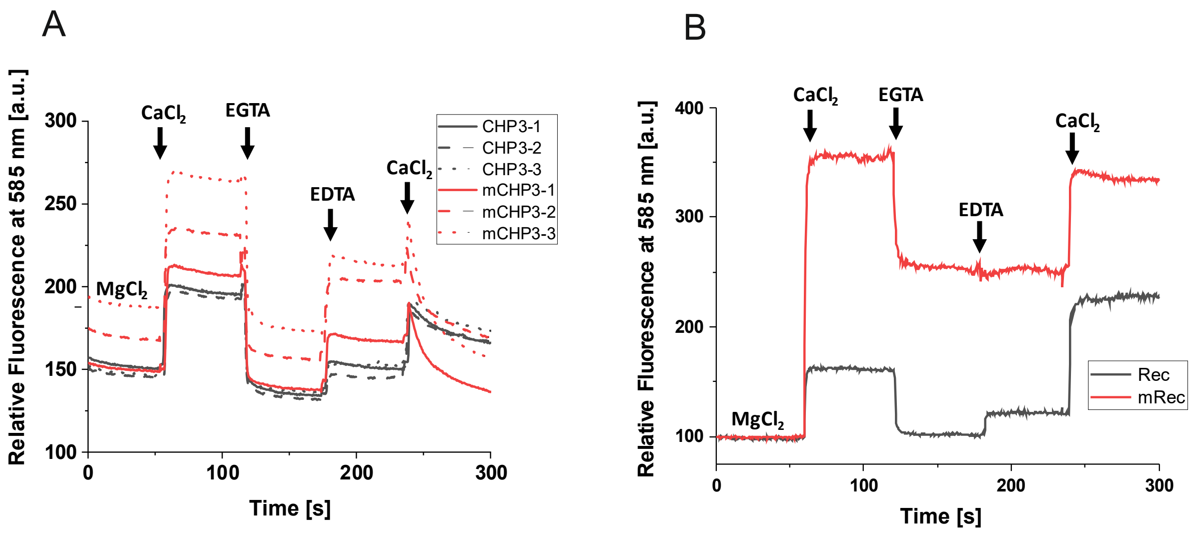


**Figure S3** Conformational changes of CHP3, mCHP3, recoverin and myristoylated recoverin monitored with the FPH assay. (**A**) Original fluorescence traces of the FPH assay for three biological replicates of CHP3 and mCHP3; (**B**) Normalized fluorescence traces of the FPH assay for myristoylated and non-myristoylated recoverin (mRec and Rec).

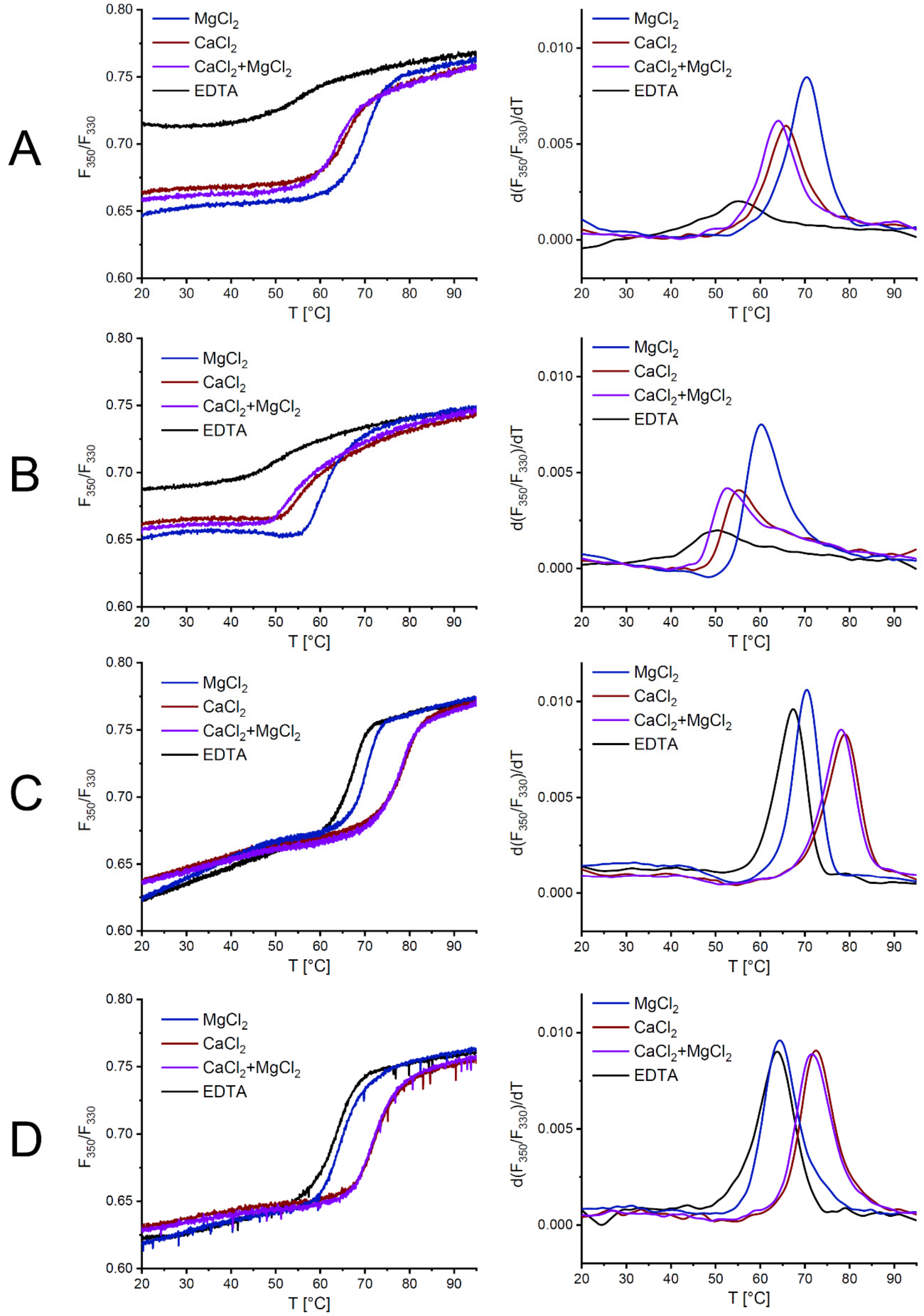


**Figure S4** Thermal unfolding of CHP3 (**A**), mCHP3 (**B**), CHP3:CBD (**C**), and mCHP3:CBD (**D**) shown as raw nanoDSF traces (left) and first derivatives (right) in the presence of either $Mg^{2+}$, $Ca^{2+}$, $Mg^{2+}$+$Ca^{2+}$, or in the absence of both ions (EDTA).

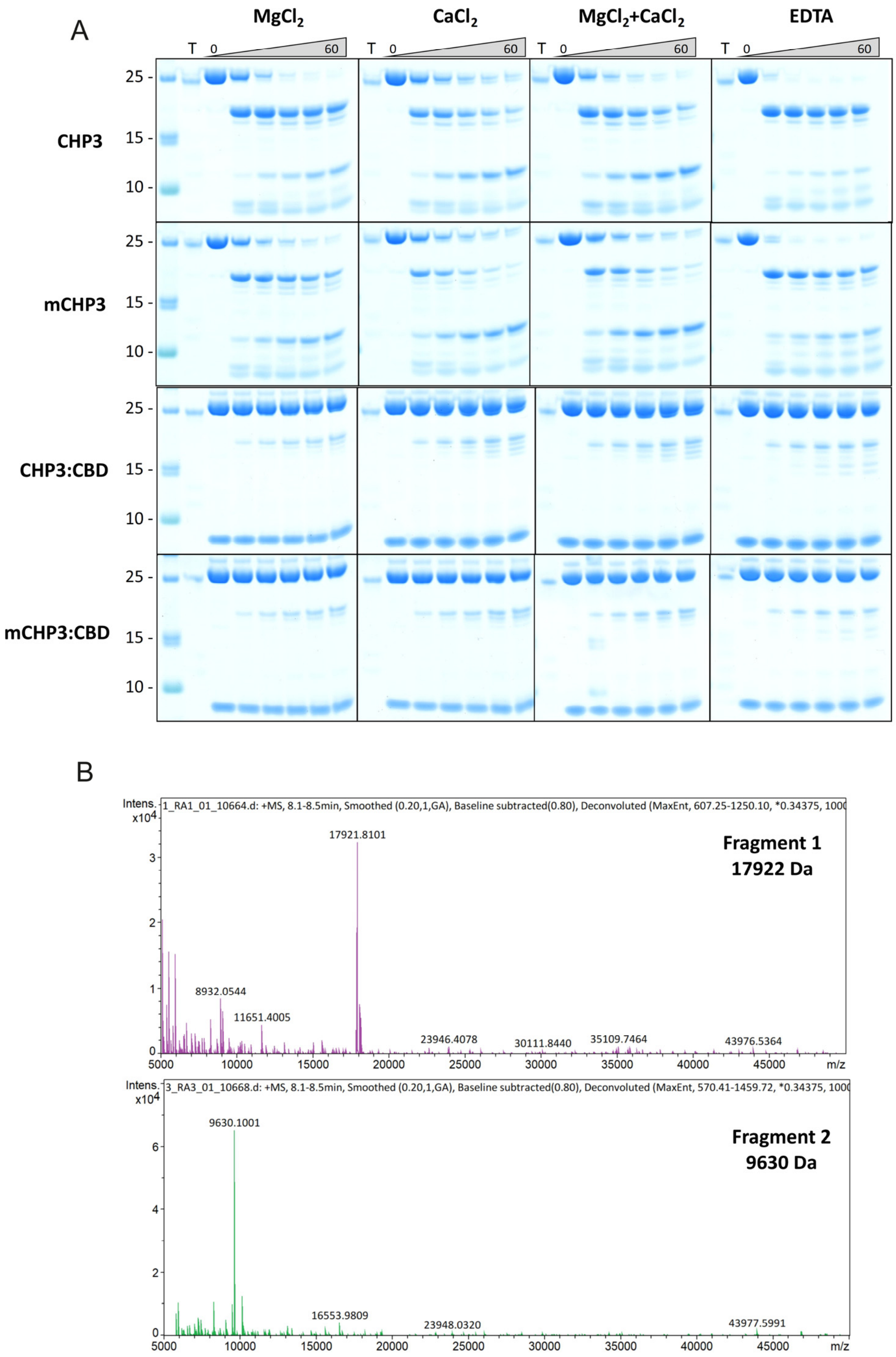


**Figure S5** (**A**) The full-size gels of limited proteolysis in different conditions. Proteins samples are indicated on the left, the ion conditions – on the top, trypsin only (T), or different CHP3 samples (CHP3, mCHP3, CHP3:CBD, or mCHP3:CBD) incubated with trypsin for 0, 5, 15, 30, 45, and 60 min were loaded from the left to the right. For a band annotation, see Figure 4A. (**B**) LC-ESI-TOF mass-spectrometry analysis of major proteolytic fragments (1 and 2), the detected masses were 17922 Da for the fragment 1 and 9630 Da for the fragment 2.

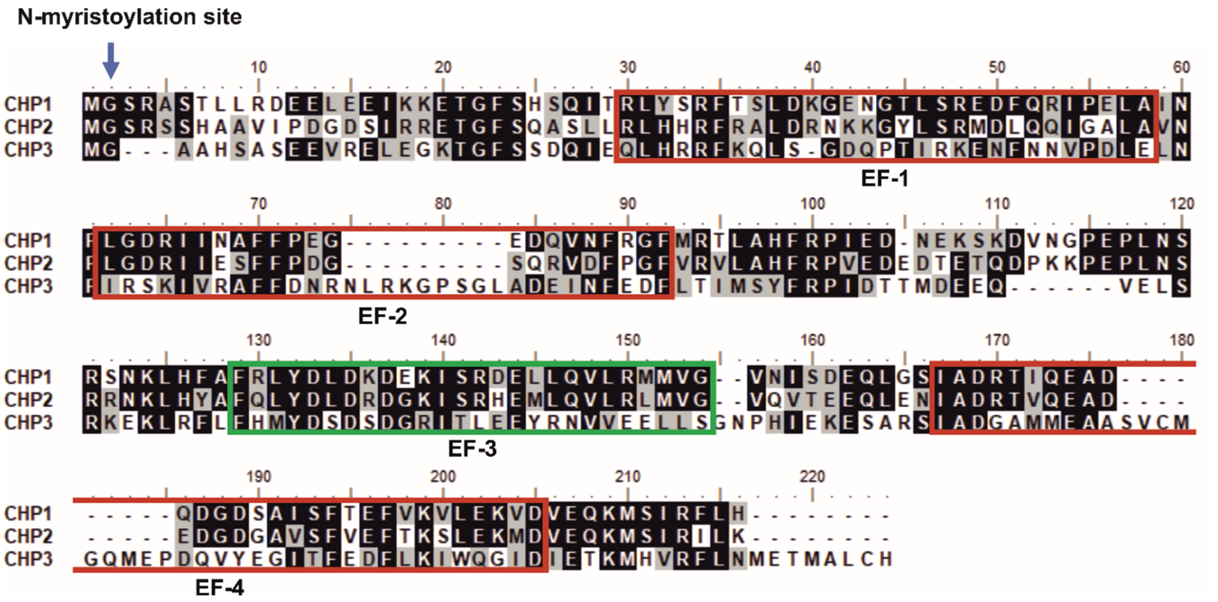

**Figure S6 Multiple sequence alignment of human CHP1, CHP2, and CHP3.** EF-1, EF-2 and EF-4 that do not bind $Ca^{2+}$ in CHP3 are shown in red rectangles, the active CHP3 EF-3 is in green, N-terminal myristoylation site is indicated with an arrow. Sequence alignment was done in BioEdit (1) using ClustalW algorithm.

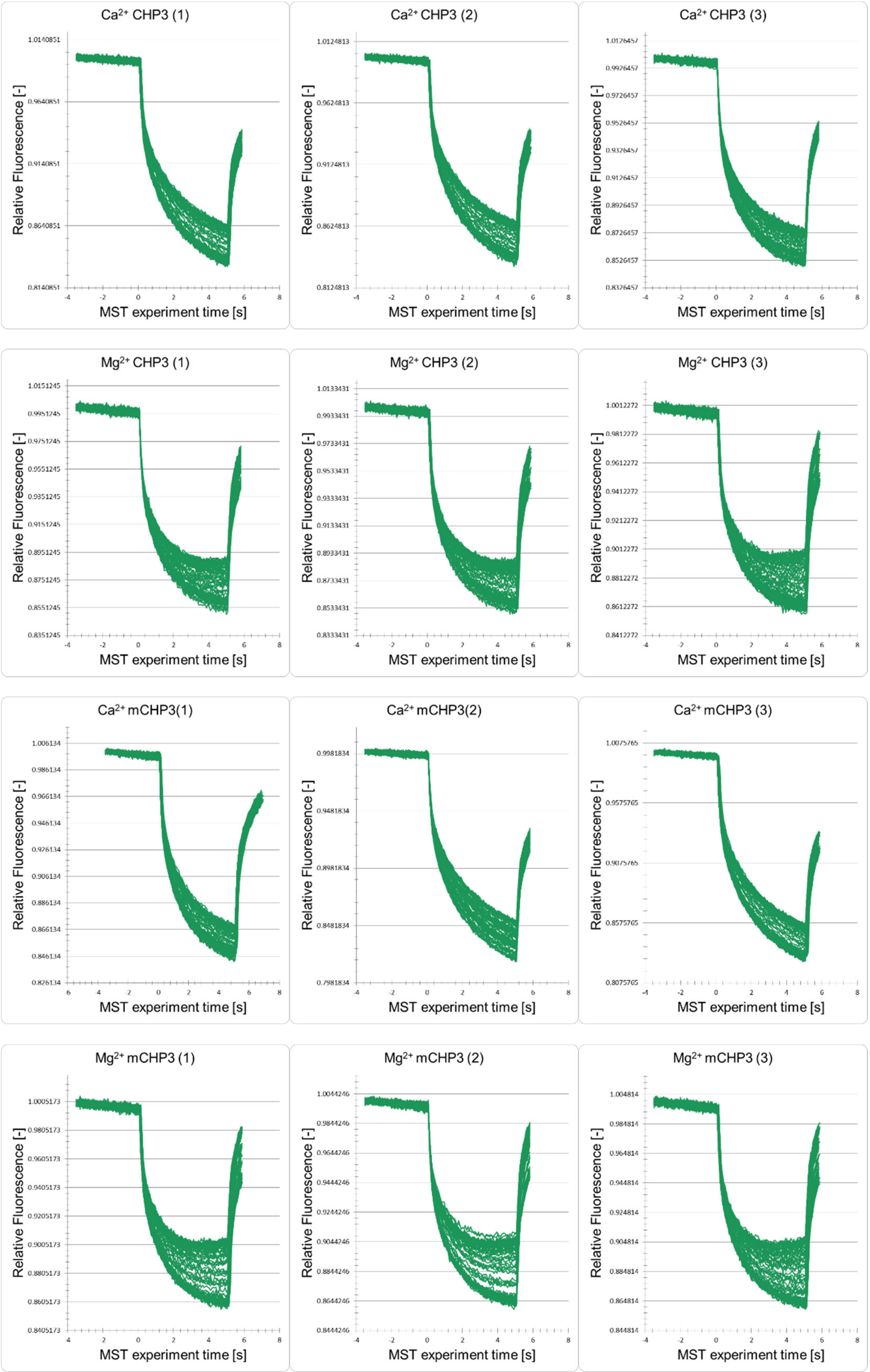


**Figure S7 Normalized MST fluorescence timetraces of CHP3 and mCHP3 with MBP-CBD in the presence of either $Mg^{2+}$ or $Ca^{2+}$.** Each graph contains measurements of one biological replicate as indicated (1-3) with five technical replicates.

```
  1 MGAAHSASEE VRELEGKTGF SSDQIEQLHR RFKQLSGDQP TIRKENFNNV PDLELNPIRS KIVRAFFDNR NLRKGSSGLA
 81 DEINFEDFLT IMSYFRPIDT TLGEEQVELS RKEKLKFLFH MYDSDSDGRI TLEEYRNVVE ELLSGNPHIE KESARSIADG
161 AMMEAASVCV GQMEPDQVYE GITFEDFLKI WQGIDIETKM HIRFLNMETI ALCH
```

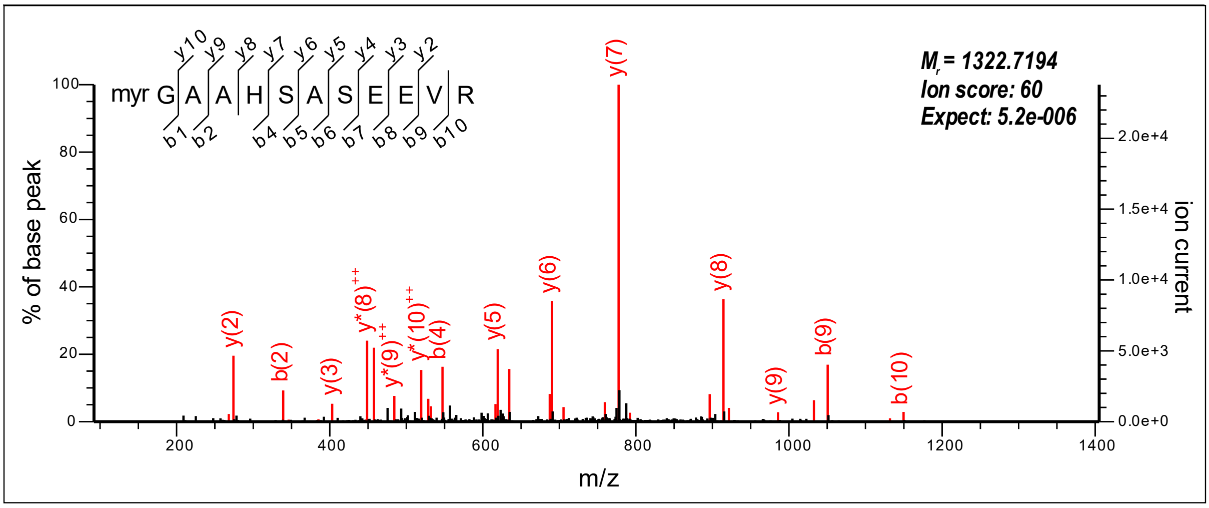


**Figure S8 Sequence coverage and N-terminal myristoylation of endogenous CHP3.** LC-MS/MS analysis of trypsin-digested total membrane fraction obtained from mouse platelets. Upper panel: Coverage of the mouse CHP3 primary sequence by MS-identified peptides (in red; black: sequences not identified, grey: sequences not accessible to our MS analysis) without inclusion of N-myristoyl modification. Lower panel: MS/MS spectrum from the same measurement assigned to the myristoylated tryptic N-terminal peptide of CHP3. No other modifications of the N-terminus were detectable in error-tolerant search.
